# Cancer-associated fibroblasts drive metabolic heterogeneity in KRAS-mutant colorectal cancer cells

**DOI:** 10.1101/2025.09.30.679631

**Authors:** Elizabeth Elton, Niki Tavakoli, Handan Cetin, Stacey D. Finley

**Affiliations:** Alfred E. Mann Department of Biomedical Engineering, University of Southern California, Los Angeles, 90089, California, USA; Department of Quantitative and Computational Biology, University of Southern California, Los Angeles, 90089, California, USA; Mork Family Department of Chemical Engineering and Materials Science, University of Southern California, Los Angeles, 90089, California, USA

## Abstract

KRAS-mutant colorectal cancer (CRC) is characterized by metabolic reprogramming that can lead to tumor progression and drug resistance. The tumor microenvironment (TME) plays a pivotal role in modulating these metabolic adaptations. In particular, cancer-associated fibroblasts (CAFs), which make up a large portion of the TME, have been shown to strongly contribute to metabolic reprogramming in CRC. This study applies flux sampling, a computational method that explores the full range of feasible metabolic states, combined with representation learning and hierarchical clustering, to a computational model of central carbon metabolism to understand how CAFs influence metabolic adaptations of KRAS-mutant CRC cells following targeted enzyme knockdowns. Focusing on twelve key enzymes involved in glycolysis and the pentose phosphate pathway, knockdowns were simulated under both normal CRC media and CAF-conditioned media (CCM) conditions. Analysis revealed that CCM induces greater metabolic heterogeneity, with knockdown models exhibiting more variable and distinct metabolic states compared to those cultured in normal CRC media. While some enzyme knockdowns produced similar metabolic states, this overlap was less frequent in CCM, indicating that CAF-derived factors diversify the metabolic responses of CRC cells to enzyme perturbations. Pathway-level flux analysis demonstrated media-specific shifts in central carbon metabolism pathways. Importantly, the predicted biomass flux showed that enzyme knockdowns reduced growth across both conditions, but models in the CCM condition indicated CAFs could offer a protective effect against metabolic perturbation. Overall, this study reveals that CCM significantly influences the metabolic state and adaptability of KRAS-mutant CRC cells to enzyme perturbations, emphasizing the importance of including TME components in metabolic modeling and therapeutic development. These findings provide valuable insights into the metabolic adaptability of CRC and suggest that targeting tumor-CAF metabolic interactions may improve treatment strategies.

**Graphical Abstract:** 

**Overview of computational workflow:** Models of interest represent simulated enzyme knockdowns in central carbon metabolism. Flux sampling searches the entire metabolic solution space and results in a distribution of flux values for each reaction within each model. Samples can be organized by knockdown and condition into matrices for input into representation learning. Representation learning is applied to sampling data to identify shared and independent metabolic states. Metabolic states indicate a heterogeneous response to enzyme knockdowns. Overlap of dark and light blue flux distributions, sampling clusters, and metabolic responses exemplify a shared metabolic state separate from to the gray unperturbed state. This workflow provides a low-dimensional representation of metabolic state that captures both the pathway- and reaction-level differences that describe each simulated knockdown.

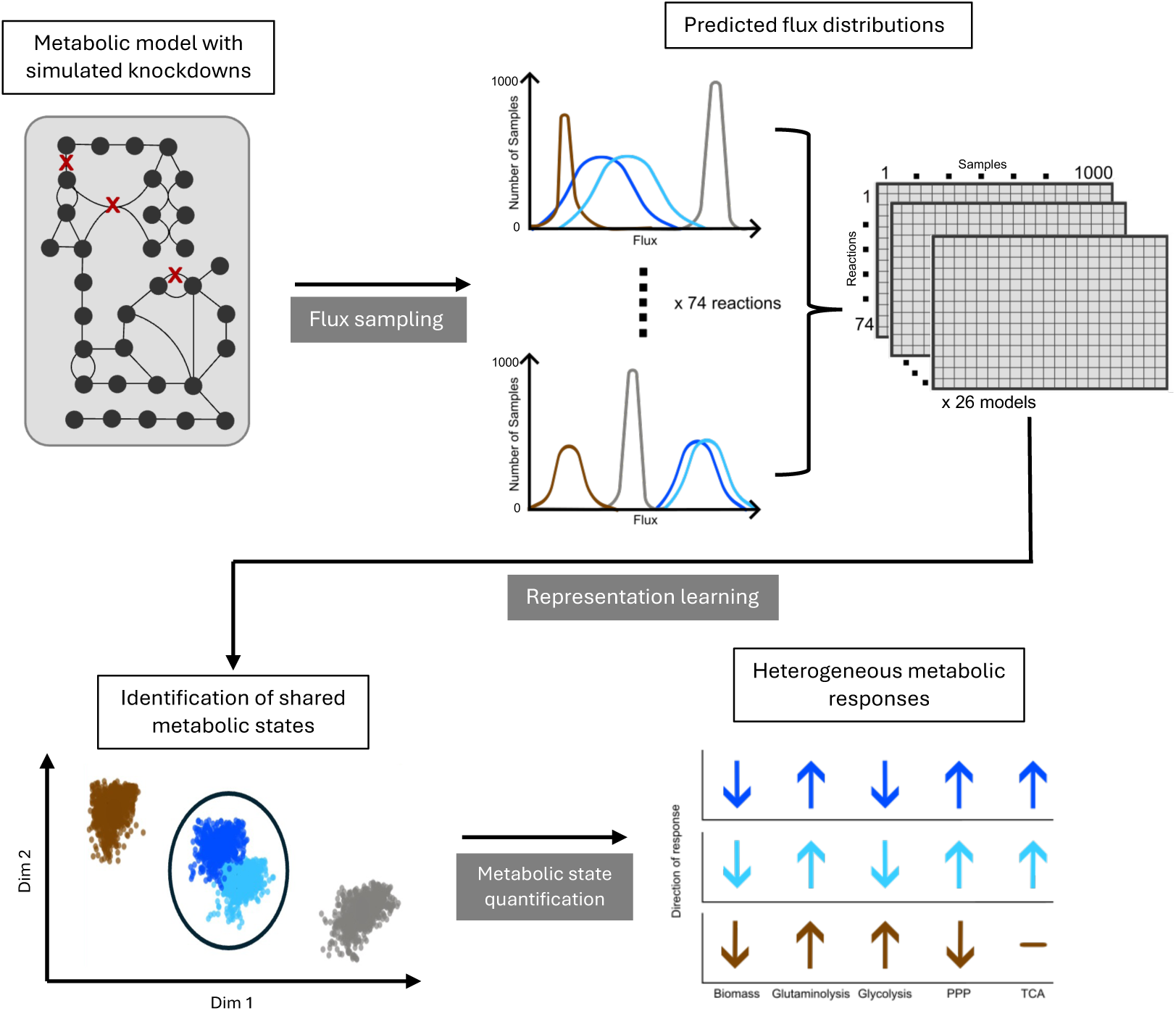

## Introduction

Colorectal cancer (CRC) is the third most common cause of cancer-related mortality worldwide, with approximately half of all CRC cases characterized by a KRAS mutation ^1,2^. In CRC, KRAS mutations drive increased cell proliferation, metastasis, and resistance to therapy^3^. These mutations are notoriously difficult to target and often reduce the efficacy of other therapies^4^. Consequently, KRAS-mutant CRC presents fewer therapeutic options, underscoring the urgent need to identify novel drug targets.

KRAS mutations are also correlated to altered cancer cell metabolism, leading to increased nutrient uptake and synthesis of essential biomolecules^5^. Given these observations, cancer metabolism has emerged as a promising area for therapeutic intervention. Metabolic reprogramming is a hallmark of cancer cells, in which these cells preferentially utilize distinct metabolic pathways compared to their normal counterparts^6^. A well-known example is the Warburg effect, where cancer cells favor aerobic glycolysis over oxidative phosphorylation for energy production, even in the presence of oxygen^7^. Effectively targeting cancer metabolism requires a deep understanding of metabolic pathways and how they are altered through cell-cell interactions within the tumor system. Although several metabolism-targeting drugs are currently in development, environmental factors within the tumor microenvironment (TME) can significantly impact therapeutic efficacy^8^. Thus, components of the TME should be considered when evaluating the effects of anti-cancer strategies that target cellular metabolism.

Cancer cell metabolism is dependent upon the availability of extracellular metabolites, which are influenced by the TME, a complex system composed of tumor, immune, and stromal cells, including fibroblasts. Cancer-associated fibroblasts (CAFs) are activated fibroblasts that support tumor progression through extracellular matrix remodeling, immune evasion, and promoting metabolic reprogramming in neighboring cells^9^. CAFs are metabolically active, and their interactions with tumor cells reshape the metabolism of both populations. For instance, in the reverse-Warburg effect, CAFs primarily utilize glycolysis to produce lactate, which is then taken up by tumor cells to fuel oxidative phosphorylation^10^. Given CRC’s high proliferative rate and its well-documented stromal-tumor interactions, the metabolic crosstalk within the TME represents a compelling target for therapeutic development. In particular, glycolytic enzymes are currently under investigation as drug targets^11^. However, a deeper understanding of the complex interactions between heterogeneous CAF populations and CRC cells is essential to exploit these metabolic interactions and predict treatment responses.

Experimental methods for studying cancer metabolism can be limited in their ability to fully recapitulate the *in vivo* environment, resource intensive, or have an unintended physiological impact. For example, isotope tracing methods can be used to quantify metabolites in biological samples yet they rely on the assumption that metabolism is at steady state, and the isotope itself can impact metabolism^12,13^. To understand how individual reactions influence cellular metabolism, a systems biology approach is necessary. This approach integrates experimental data with computational models to study the interactions between components of a biological system as a whole. In cancer metabolism, systems biology aims to quantitatively describe the utilization of all metabolic reactions within cells, particularly in the TME. Given the limitations of experimental methods alone, computational tools are essential for predicting cellular metabolic behavior and identifying novel therapeutic strategies, particularly in CRC, where intercellular metabolic interactions drive disease progression.

A computational systems biology framework has already provided meaningful biological insights into cancer metabolism. In particular, constraint-based modeling allows for the prediction of reaction fluxes and cell growth rates. Wang et al. used this approach to integrate metabolomics data into a CRC-specific model of central carbon metabolism^14^. They developed condition-specific models of KRAS-mutant and wild-type CRC cultured with or without CAFs and predicted how knockout of individual metabolic enzymes affected tumor cell growth and response to CAF-conditioned media. The study demonstrated that CAFs rewire CRC metabolism by increasing flux through glycolysis, the oxidative pentose phosphate pathway (PPP), and glutaminolysis, while reducing activity in the TCA cycle. Tavakoli et al. expanded this work by simulating partial knockdowns of metabolic enzymes to predict effects on a broader set of central carbon metabolic reactions^15^. Their model predicted that inhibiting HK would have distinct effects depending on whether CRC cells were cultured in CAF-conditioned media (CCM) versus standard CRC media, and these predictions were validated experimentally. Both works utilized flux balance analysis (FBA) to draw their conclusions, where FBA is a computational method to find the flux through each reaction in a metabolic model given a set of constraints and a measured growth rate^16^. FBA, however, is limited in that it only provides one optimized solution set of predicted flux values – namely, the flux values that optimize a user-defined objective function. When studying cancer cell metabolism, this objective function is usually to maximize growth while minimizing overall energy use^15,17^. However, after implementing a perturbation that influences intracellular metabolism, this assumption that the cell still optimizes growth may not hold true. We must instead explore all possible flux values for each reaction to understand the variety of metabolic adaptations taken by CRC cells^18,19^. Unlike traditional FBA, flux sampling captures the full range of possible pathway-specific flux distributions under given constraints. We therefore aim to use flux sampling to expand upon an existing model of central carbon metabolism to understand the range of metabolic responses CRC cells can undergo following a metabolic perturbation.

Flux sampling has been leveraged in similar contexts, including exploring metabolic heterogeneity in immune cells within CRC. For example, flux sampling was used to compare M1 and M2 macrophage metabolic states, revealing key reactions that distinguish these phenotypes^20^. Flux sampling has also proven effective in identifying targetable reactions and characterizing metabolic variability; Piccialli et al. demonstrated that clustering analyses of flux sampling results can distinguish metabolic states across different cancer cell lines using publicly available datasets^21^.

Computational modeling thus offers a powerful platform to explore how metabolic states emerge in response to specific cellular objectives. However, many existing studies focus on optimized or steady-state metabolic conditions, and relatively few have explored how metabolic heterogeneity arises following targeted perturbations of metabolic pathways, particularly in a cell-diverse environment like the TME^22,23,24^. While these works offer helpful insights into our understanding of cancer metabolism, they introduce the question: how can we understand the metabolic adaptations that are undertaken by cancer cells in a mixed-cell environment after perturbation? This work, therefore, seeks to investigate the metabolic heterogeneity of CRC cells in the context of the TME after targeted enzymatic perturbation, as these perturbations can lead to diverse metabolic responses and uncover therapeutic vulnerabilities that are not evident under optimal conditions. We hypothesize that flux sampling will provide new insight into the metabolic states present in CRC cells cultured in normal media compared to CRC cells cultured in CCM.

Here, we utilized flux sampling to investigate the range of possible metabolic states that arise from targeted perturbations of a model of central carbon metabolism in KRAS-mutant CRC cells. Specifically, we examined how this metabolic heterogeneity is shaped by the influence from CAFs in the TME. We applied flux sampling to constraint-based models of metabolism informed by previously performed experiments using two media conditions: CRC cells cultured in their own media and those cultured in CAF-conditioned media, which describes growing CRC cells in media that CAFs had previously grown in^14^. The base (unperturbed) model consists of 74 reactions representing central carbon metabolism, including a biomass reaction, acting as a proxy for cellular biomass growth. We simulated enzyme perturbations identified in prior work by Tavakoli et al. as having a significant impact on CRC metabolism. Flux sampling using the base and enzyme knockdown models reveal that each media condition is associated with distinct metabolic states; CRC cells grown in CAF-conditioned media exhibited greater metabolic heterogeneity, as reflected by pathway flux distributions and predicted biomass production. These results highlight the influence of CAF-tumor interactions in shaping metabolic responses to perturbation and underscore the need to consider the TME when designing metabolism-targeting therapies. Furthermore, our work contributes to the identification of metabolic vulnerabilities for a resistant cancer type (KRAS-mutant) and offers a flexible computational workflow that can be adapted for broader studies of metabolic crosstalk. By investigating the full spectrum of metabolic states that emerge from targeted enzyme inhibition, this study provides new insights into CRC metabolism and supports the development of effective strategies for metabolic intervention in cancer.

## Results

### Flux sampling reveals the range of reaction fluxes in the metabolic model of central carbon metabolism in CRC cells

In prior work from our research group, constraint-based modeling was used to simulate the effect of enzyme perturbations in central carbon metabolism in CRC cells on biomass^14^. Building on that analysis, Tavakoli et al. identified impactful metabolic perturbations that influence central carbon metabolism, rather than focusing solely on biomass production^15^. In the most recent work, machine learning was used to quantify the effects of enzyme perturbations predicted by a computational model of CRC metabolism. Specifically, representation learning casts the output of the computational metabolic models into 2-D space^15^. In this representation, the distance between any two points reflects the degree of similarity between their corresponding metabolic states, where smaller distances indicate more similar states. This approach allows for comparison of the effects of individual enzyme knockdowns both within a particular media condition and across different media conditions. This resulted in 12 enzyme knockdowns, including 11 knockdowns involved in glycolysis, and one knockdown in the pentose phosphate pathway (PPP). These enzyme knockdowns are defined in **Table 1**. Some enzymes knockdowns correspond to the same enzyme but inhibited by a different percentage. For example, in one model, hexokinase (HK) is reduced by 80% of the maximum flux through the HK reaction achieved in the base model. This compares to a 100% knockdown of HK, in which zero flux can move through the reaction mediated by HK. Each of the 12 influential enzyme knockdowns were simulated for models representing two cell culture conditions: KRAS-mut cells cultured in CAF-conditioned media (CCM) and KRAS-mutant cells in CRC media. In total, including both CCM and CRC media conditions, the knockdown models, and the base model, 26 models of interest were investigated.

**Table 1.** Description of enzyme knockdowns.

| Enzyme Abbreviation | Knockdown Level Simulated (%) | Full Enzyme Name | Pathway |
| --- | --- | --- | --- |
| ALDO | 60 | Aldolase | Glycolysis |
| ENO | 100 | Enolase | Glycolysis |
| G6PDH | 100 | Glucose 6-phosphate dehydrogenase | Pentose Phosphate Pathway |
| GAPDH | 60 | Glyceraldehyde phosphate dehydrogenase | Glycolysis |
| HK | 80, 100 | Hexokinase | Glycolysis |
| HPI | 80, 100 | Glucose 6-phosphate isomerase | Glycolysis |
| LDH | 20, 40 | Lactate dehydrogenase | Glycolysis |
| PGAM | 100 | Phosphoglycerate mutase | Glycolysis |
| PYK | 100 | Pyruvate kinase | Glycolysis |

To understand the possible metabolic states of KRAS-mut CRC cells after enzyme knockdown, flux sampling was employed to capture the range of flux values through each reaction. In doing so, 1000 possible flux values were obtained for each reaction (**Supplemental Figure 1**). For some reactions, flux distributions were a single value or had a standard deviation smaller than 10^-5^ mM/h and were approximated as single values. Other reactions showed a greater range of possible values, with the greatest magnitude flux achieved approximately +/-499 mM/h. The lactate dehydrogenase (LDH) 20% and LDH 40% knockdown models consistently showed the greatest flux distribution range for each reaction. As each reaction has a distribution of 1000 possible flux values per model, we can represent each model as the 1000 solution vectors of flux values that result from sampling, where each vector is composed of one flux value for each of the 74 reactions. These solution vectors were further analyzed to define and describe the metabolic states present in each condition.

### Representation learning applied to flux sampling identifies metabolic states

Flux sampling analysis revealed that enzyme knockdowns result in specific metabolic states, which can be understood as distinct distributions of fluxes across the metabolic network. To visualize and compare these states, representation learning was utilized to project the predicted metabolic states into a 2-D space.

Inter-cluster heterogeneity was assessed by measuring the Euclidean distance between the centroid of the cluster for each model and the centroid of either the base model (within a condition) or the corresponding model under the alternate media condition. Results showed that, for any given enzyme knockdown, the metabolic states predicted for CCM and CRC conditions were clearly distinct, with no overlap observed between knockdown models across the two media types (**Figure 1A**). Additionally, knockdown models under CCM were consistently farther from the CCM base model than the CRC knockdown models were from the CRC base model. In both conditions, the PYK 100% model was the most distant from the respective base model. Interestingly, many CRC knockdown models were closer to the CRC base model than to their CCM counterparts. For example, PGAM 100% CRC was closer to the CRC base model than to PGAM 100% CCM, suggesting media condition-specific metabolic responses (**Supplemental Table 1**).

**Figure. 1.**
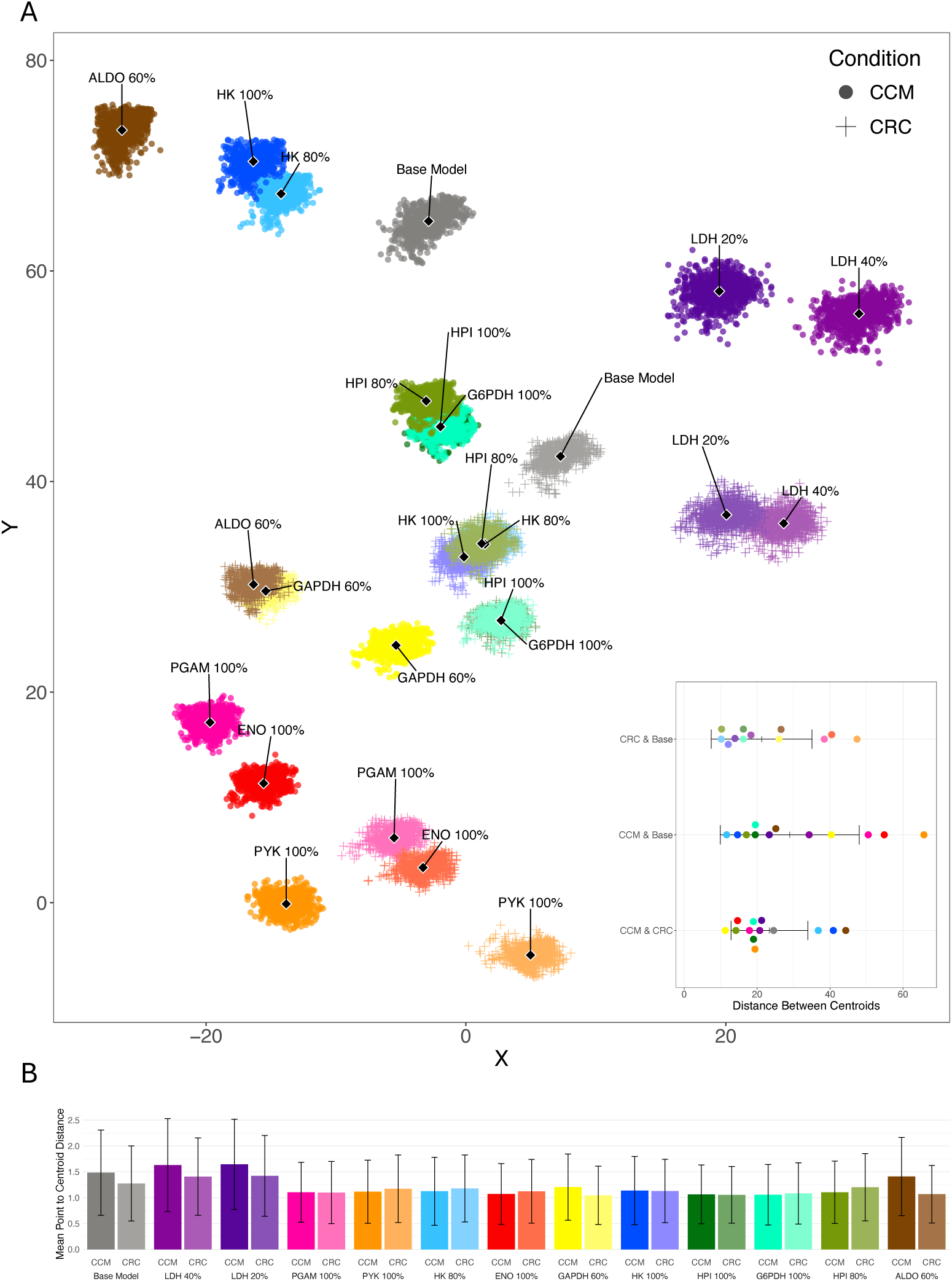
Representation learning of flux sampling data identifies metabolic states for CCM and CRC conditions. (A) Samples from all models in both conditions are projected into 2-D space using a representation learning approach. Enzyme knockdown models are represented as different colors. Each point represents one sample, which is one possible solution vector of flux values. The centroid of each model cluster is represented as a black diamond. Circles, CCM model samples; crosses, CRC model samples. The inset describes distance between model cluster centroids. Black bars indicate mean and standard deviation. (B) Spread within a cluster is determined by the distance between the cluster centroid and each point in the cluster. Black bars indicate standard deviation.

To assess intra-cluster heterogeneity, the spread of each model cluster was examined by calculating the mean Euclidean distance from each sampling point to its cluster centroid. These results reflect how variable the predicted metabolic states are within each model (**Figure 1B**). Notably, in the CCM condition, the spread from the centroid was greater for several models compared to the same knockdown in CRC media, including the Base Model, HK 80%, HK 100%, HPI 100%, G6PDH 100%, and HPI 80%. This indicates that these knockdown models exhibit greater heterogeneity in their potential metabolic states under CCM compared to their CRC counterparts.

Our results show that different enzyme knockdowns can converge to shared metabolic states. To quantify this, the percent overlap between representation learning sampling clusters was calculated. This was done by determining the convex hull for each model cluster, calculating the area of intersection between pairs of hulls, and dividing this by the non-intersection area. Shared metabolic states are defined by the percent overlap between model clusters (**Supplemental Figure 2**). Shared metabolic states can be described as having low (1-25%), intermediate (26-75%), or high overlap (76-100%). In the CCM condition, overlapping clusters, and therefore shared metabolic states, were identified among several models. An intermediate overlap was observed between the HK 80% and HK 100% knockdown models. A second state comprised of the HPI 80%, HPI 100%, and G6PDH 100% models showed stronger similarity: HPI 100% and G6PDH 100% showed high overlap, while intermediate overlap was observed between HPI 100% and HPI 80%, and HPI 80% and G6PDH 100%. In the CRC condition, shared metabolic states were observed among distinct model groups. HK 80%, HK 100%, and HPI 80% knockdown models formed one shared state, with intermediate overlap between HK 80% and HK 100%, and HK 100% and HPI 80%, and high overlap between HK 80% and HPI 80%. High overlap was also observed for GAPDH 60% and ALDO 60%, and HPI 100% and G6PDH 100% knockdown models. Intermediate overlap was observed for LDH 20% and LDH 40% and low overlap was observed for PGAM 100% and ENO 100% knockdown models. Overall, fewer shared metabolic states were found among distinct enzyme knockdowns in the CCM condition compared to CRC, suggesting that cells cultured in CCM exhibit a more heterogeneous response to metabolic perturbations.

### Clustering analysis of flux sampling data

Hierarchical clustering further supports the metabolic state distinctions identified through representation learning. To explore the similarities between predicted metabolic states, hierarchical clustering was performed using the mean flux values from the sampling distribution of each reaction, applying Euclidean distance as the similarity metric and using complete linkage as the agglomeration method (**Figure 2**). This analysis demonstrated that the magnitude of flux changes between enzyme knockdown models and the base model was greater under CCM conditions than under CRC, reinforcing earlier findings that CCM induces greater heterogeneity in metabolic responses.

**Figure 2.**
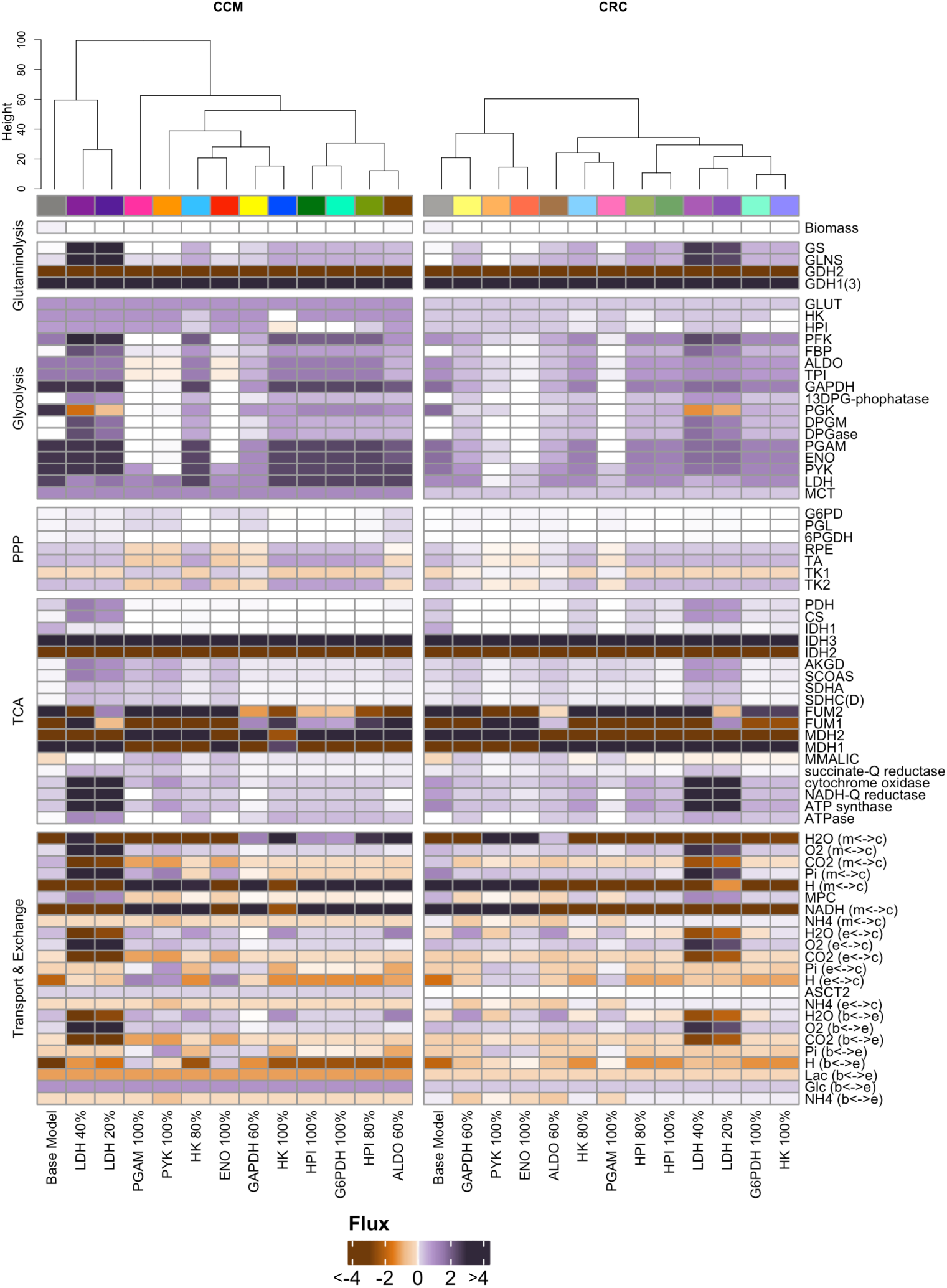
Clustering of mean flux values validates identified metabolic states. Heatmaps of mean flux through each reaction for each model under the CCM and CRC conditions. Models are indicated by color and clustered based on Euclidean distance. Reactions are grouped by metabolic pathway. The color scale illustrates flux through each reaction, with hue indicating flux values from -4 mM/h to 4 mM/h. Flux values greater than 4 mM/h or less than -4 mM/h are set at the maximum color intensity limits. Sorted dendrogram height at node indicates level of similarity between models.

The hierarchical clustering results aligned with patterns observed in representation learning, providing additional evidence of increased heterogeneity under CCM conditions. The dendrogram generated for the CCM condition displayed a greater maximum height at the top node compared to the CRC dendrogram, indicating a higher degree of overall dissimilarity between models. Additionally, in the CCM dendrogram, two models (PGAM 100% and the Base Model) were represented as separate, isolated branches beyond the second node, while no isolated branches were present in the CRC dendrogram, suggesting a broader distribution of metabolic states in CCM.

Several shared metabolic states identified through representation learning were also recapitulated in hierarchical clustering. For example, in CCM, the shared metabolic state formed by HPI 80%, HPI 100%, and G6PDH 100% knockdowns clustered closely together. This group was distinguished by increased flux through glycolytic reactions (GAPDH, diphosphoglycerate mutase (DPGM), and diphosphoglycerate phosphatase (DPGase)), decreased flux through PPP reactions (G6PDH, phosphogluconolactonase (PGL), and 6-phosphogluconate dehydrogenase (6PGDH)), and similar flux values in transport and exchange reactions involving CO₂, Pi, and H⁺. Notably, the HPI 100% and G6PDH 100% models showed the highest percent overlap in representation learning (**Figure 1A**) and exhibited nearly identical flux values across all reactions.

Similarly, in the CRC condition, the shared metabolic state formed by LDH 20% and LDH 40% knockdowns was also evident in the hierarchical clustering analysis. These two models exhibited nearly identical flux values across most reactions, apart from FUM1 and FUM2, which showed a negative flux in the LDH 20% model and a positive flux in the LDH 40% model.

Beyond clustered groups, some knockdown models that did not form shared clusters in either representation learning or hierarchical clustering still demonstrated common trends in predicted flux profiles. These patterns were especially pronounced in the CCM condition, where the PGAM 100%, ENO 100%, and PYK 100% models all predicted decreased flux through multiple glycolytic reactions, including negative flux through ALDO and TPI. The models also showed negative flux through PPP reactions such as RPE, TA, and TK2, in contrast to the positive fluxes predicted for these reactions in all other models, including the base model.

Interestingly, while similar flux trends were observed for PGAM 100% and ENO 100% in both CCM and CRC conditions, the magnitude of change relative to the base model was significantly greater in CCM. For example, these models predicted negative flux through RPE, TA, and TK2 under both conditions, but the absolute values of these fluxes were higher in CCM, further demonstrating the media-specific alteration of responses.

### Summed pathway flux describes shared metabolic states

Each metabolic state was quantitatively defined by analyzing the flux through central carbon metabolism pathways. This was achieved by summing the flux through each pathway per sampling instance and calculating the mean pathway-level flux for each model under both CCM and CRC conditions (**Figure 3**). Through this analysis, common trends in pathway-level fluxes were identified among enzyme knockdown models with shared metabolic states, as determined by representation learning. Additionally, consistent flux patterns were observed across certain models that, while not grouping together in representation learning, still demonstrated notable similarities at the pathway level.

**Figure 3.**
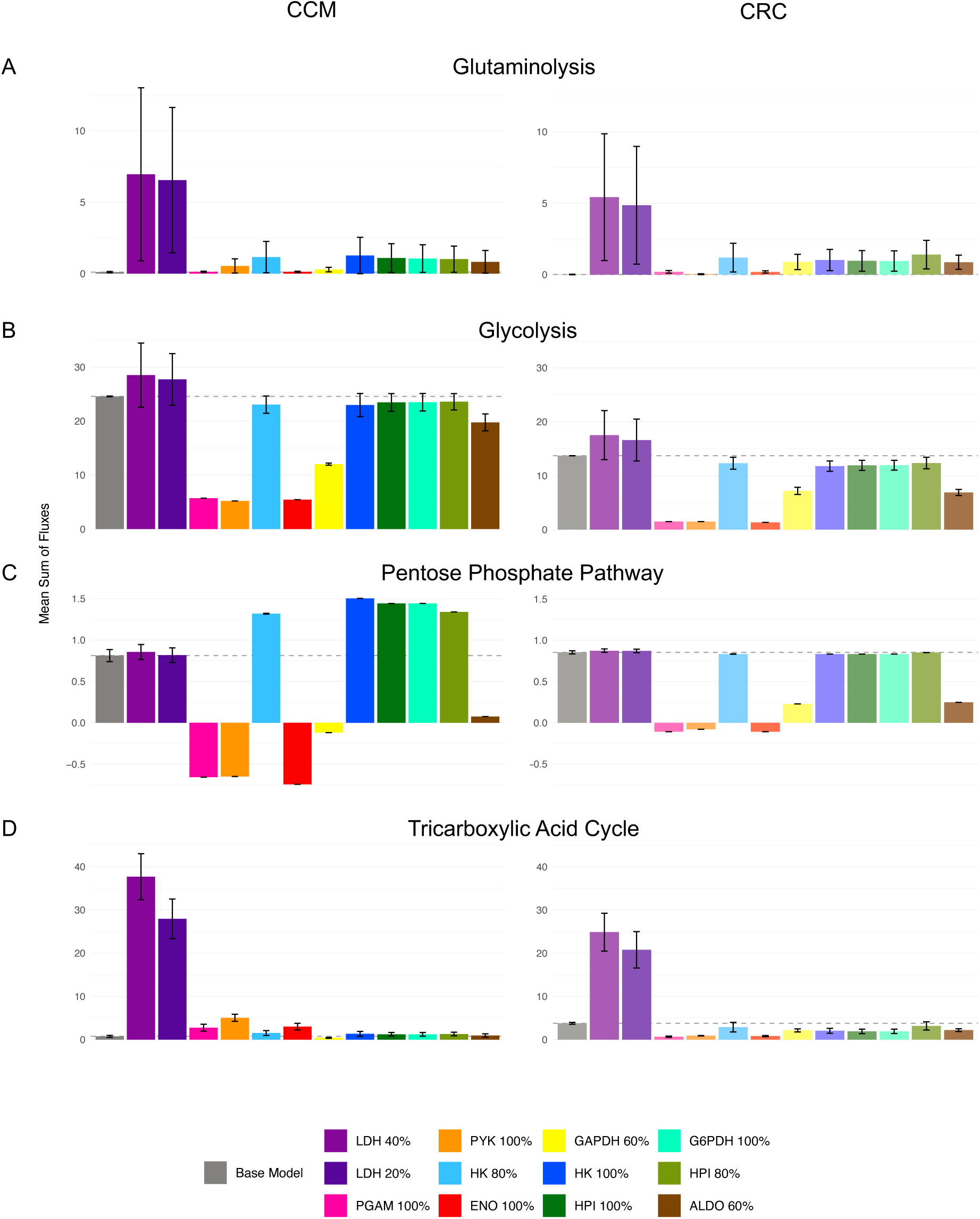
Pathway flux describes shared metabolic states. Mean of the sum flux (mM/h) through (A) glutaminolysis, (B) glycolysis, (C) pentose phosphate pathway, and (D) tricarboxylic acid cycle for each model under the CCM condition (left) and the CRC condition (right). For each sample, the flux through each reaction is grouped by pathway and added. The sum flux through the Base Model is indicated by gray dashed line. Error bars indicate standard deviation.

In the case of glutaminolysis (**Figure 3A**), both CCM and CRC base models showed near-zero or zero predicted flux. Most knockdown models in both conditions displayed increased or similar flux through glutaminolysis relative to baseline. Under CCM, the HPI 100%, HPI 80%, and G6PDH 100% models predicted glutaminolysis flux levels similar to those of the HK 80% and 100% knockdown models. This shared pattern contributes to defining the metabolic state associated with the HPI and G6PDH knockdowns in CCM, indicating that glutaminolysis flux can help characterize shared metabolic shifts.

For glycolysis (**Figure 3B**), the predicted flux changes in knockdown models were consistent across media conditions. LDH knockdown models showed increased glycolytic flux, while ENO 100%, PYK 100%, PGAM 100%, GAPDH 60%, and ALDO 60% models predicted decreased glycolysis relative to baseline. Models such as HPI 80% and 100%, HK 80% and 100%, and G6PDH 100% showed similar or slightly reduced glycolytic flux compared to the base model. Importantly, all CCM models predicted greater glycolysis flux than their CRC counterparts, suggesting enhanced glycolytic activity in the CCM condition across perturbations.

Flux through the PPP (**Figure 3C**) revealed additional distinctions between metabolic states. Four models in CCM (PGAM 100%, ENO 100%, PYK 100%, and GAPDH 60%) predicted negative flux through the PPP, indicating a reversal of the non-oxidative branch of the pathway. Notably, PGAM 100%, ENO 100%, and PYK 100% showed a greater magnitude of negative PPP flux under CCM than CRC, suggesting a shift toward the production of reactants rather than products. GAPDH 60% had positive PPP flux in CRC but negative in CCM. Base model PPP flux was comparable across conditions, but in CCM, knockdown models such as HPI 80% and 100%, G6PDH 100%, and HK 80% and 100% predicted increased PPP flux compared to baseline. These same models in CRC had PPP flux values similar to the base model.

Within the tricarboxylic acid cycle (TCA) (**Figure 3D**), the base model flux was lower in CCM than in CRC. In CCM, nearly all knockdown models predicted similar or increased TCA flux compared to the base model. In comparison, in the CRC media condition, TCA flux was similar to or decreased in all models except LDH knockdowns, which maintained or increased TCA activity relative to baseline. These differences indicate a broader metabolic response in CCM knockdowns that leads to increased TCA usage compared to the base model, despite its overall lower activity compared to the CRC media condition.

Interestingly, several models that do not cluster together in representation learning or hierarchical clustering still showed similar pathway-level flux patterns. For example, under CCM, HPI 80% and 100% and G6PDH 100% knockdown models exhibited similar flux profiles to the HK 80% and 100% models across multiple pathways. Similarly, ENO 100%, PGAM 100%, and PYK 100% knockdown models predicted comparable fluxes in glutaminolysis, glycolysis, PPP, and biomass production. The GAPDH 60% model under CCM shared similar pathway flux trends with PGAM 100%, PYK 100%, and ENO 100%, while under CRC it more closely resembled HPI 80% and 100%, G6PDH 100%, and HK 80% and 100% models. This observation may reflect the generally greater homogeneity in flux values under CRC conditions compared to CCM, reinforcing our findings of increased metabolic heterogeneity in the CCM condition described above.

### Predicted biomass flux after enzyme knockdown

The biomass reaction serves as a representation of cellular growth, connecting metabolic activity to physiological outcomes. Thus, it is useful to compare the predicted biomass flux for the knockdown models and culture conditions. Across all models and media conditions, enzyme knockdowns resulted in decreased predicted biomass flux compared to the base model (**Figure 4**). However, the extent of this decrease varied depending on both the specific enzyme knocked down and the media condition, highlighting the complex, condition-dependent nature of metabolic perturbations.

**Figure 4.**
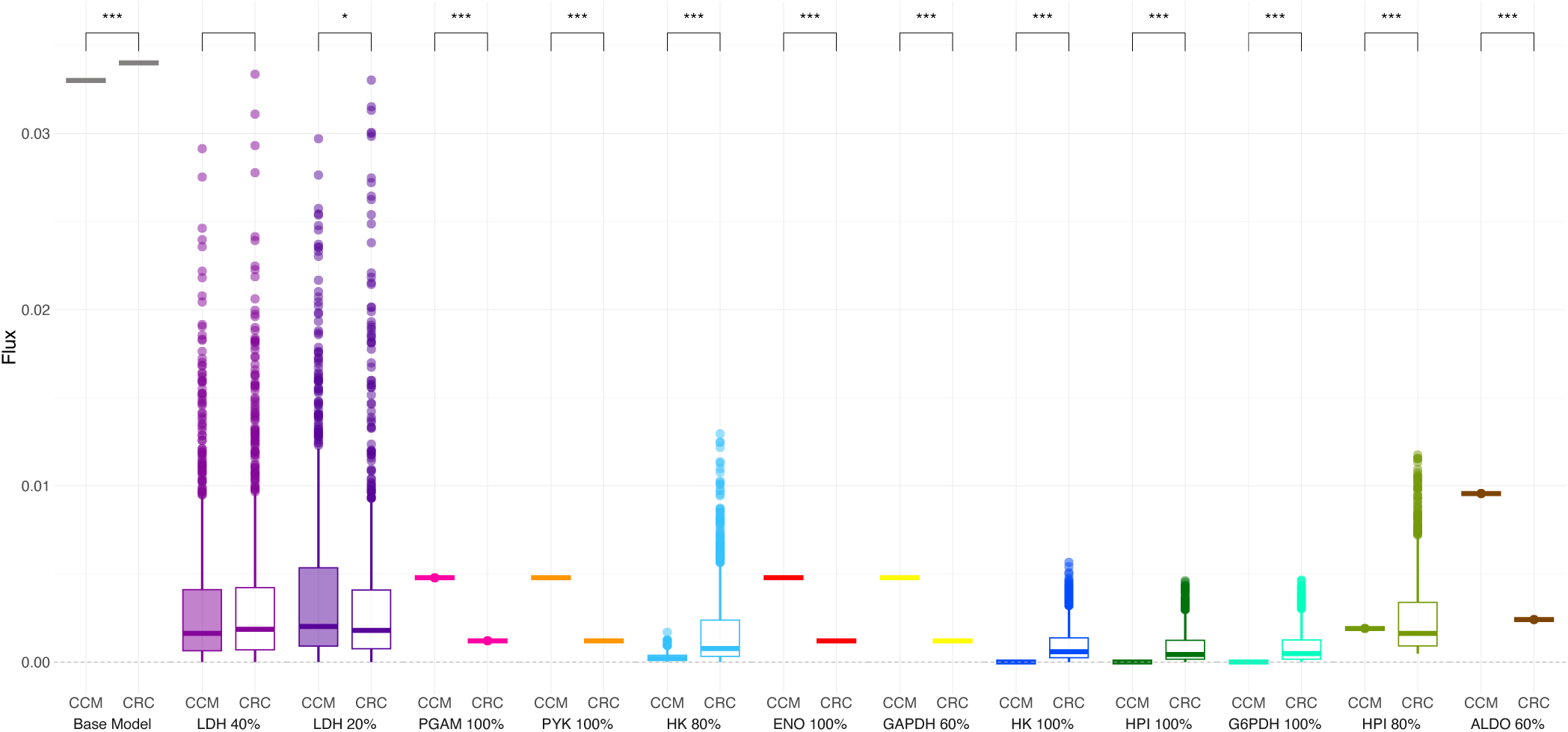
Distributions of predicted biomass flux values represent cellular growth after perturbation. Sampled flux (mM/h) through the biomass reaction for each model under the CCM condition (left, filled) and the CRC condition (right, open). Results of Kolmogorov-Smirnov test indicated above brackets (***<0.001, **<0.01, *<0.05)

There was greater variability in predicted biomass flux under CRC conditions compared to CCM. Most CCM knockdown models produced only a single value for biomass flux, suggesting a narrower range of possible growth outcomes. In contrast, CRC models exhibited a broader distribution of biomass flux values. Quantitatively, the standard deviation of mean predicted biomass flux across all knockdown models (excluding the base model) was 0.003 in CCM and 0.001 in CRC (**Supplemental Table 2**). This supports our findings that enzyme knockdowns lead to more heterogeneous metabolic states in CCM, while CRC models tend to converge toward similar outcomes, particularly in terms of growth potential.

Interestingly, some CRC models predicted lower biomass flux than their CCM counterparts. This observation suggests that models in the CCM condition are demonstrating the protective effect of CAFs on CRC cells, making them more resilient to perturbations. Statistical analysis of biomass flux distributions revealed significant differences between the CCM and CRC conditions for nearly all models (Kolmogorov-Smirnov test, FDR-corrected p-value < 0.05), except for LDH 40%, where no significant difference was detected. In terms of media-specific effects, the mean predicted biomass flux was lower in CRC for several knockdown models, including LDH 20%, PGAM 100%, PYK 100%, ENO 100%, GAPDH 60%, HPI 80%, and ALDO 60%. Conversely, the CCM condition resulted in lower biomass flux in the HK 80%, HK 100%, HPI 100%, G6PDH 100%, and base models.

Notably, several knockdown models under the CCM condition predicted zero biomass flux, indicating complete inhibition of cell growth. These included HK 100%, HPI 100%, and G6PDH 100%, while HK 80% had the next lowest predicted biomass flux, with a mean value markedly lower than the value predicted for the same model under CRC. These results suggest that certain knockdowns could lead to greater growth inhibition effects in the CCM environment.

The condition-dependent impact on biomass flux observed in HK knockdown models aligns with findings from the Tavakoli et al. study, which reported that drugs targeting hexokinase activity are more effective at disrupting cellular metabolism in CCM than in CRC. Taken together, these findings underscore the importance of incorporating the tumor microenvironment in determining the outcomes of metabolic perturbations.

## Discussion

Our results demonstrate that CCM leads to distinct metabolic states in KRAS-mutant CRC cells after simulated enzyme knockdowns. While prior studies have established that CAFs influence CRC metabolism and identified enzyme knockdowns with significant effects on optimized metabolic states in CRC, our results go beyond this conclusion to elucidate the diverse set of adaptations that CRC cells take after perturbation^14,15^. Importantly, our results robustly show that simulated knockdowns of different enzymes can lead to similar metabolic responses particular to CCM. This convergence to shared metabolic states occurs less frequently in CCM media, suggesting that the presence of factors secreted by CAFs during growth can alter the pathways CRC cells employ in response to metabolic perturbations. These findings have meaningful impacts for KRAS-mutant CRC and the identification of new treatment strategies.

The metabolic heterogeneity resulting from enzyme knockdowns in the CCM condition is indicative that CAFs alter the metabolic adaptations of KRAS-mutant CRC cells in response to perturbation. Representation learning illustrated that knockdowns of the same enzyme by different percentages could lead to different metabolic outcomes. Conversely, in both conditions, knockdowns of different enzymes led to shared metabolic states. We note the presence of fewer shared metabolic states overall in the CCM condition compared to the CRC growth condition, indicating that CCM diversifies the metabolic recovery strategies of CRC cells. Hierarchical clustering analysis leads to a similar finding: the top node in the CCM dendrogram occurred at a greater height than the CRC dendrogram, indicating less similarity between the knockdown model flux predictions. This observation is supported by prior studies demonstrating the impact of intra-tumoral metabolic heterogeneity and the known diversity of CAF populations^25,26,27^.

In many cases, CRC knockdown models remained more similar to their unperturbed CRC base model than to the corresponding knockdown in CCM. We note that our application of representation learning and hierarchical clustering produced this result, despite differences in the methods. Namely, representation learning emphasizes similarities between inputs, while clustering analysis accentuates differences between inputs^28^. For instance, representation learning showed that the PGAM knockdown in CRC media more closely resembled the CRC base model than the same knockdown in CCM. Hierarchical clustering reached a similar conclusion, where the CCM base model separated from knockdown models at a higher node in the dendrogram than the corresponding CRC models, suggesting a broader range of responses in the CCM condition.

Both representation learning and hierarchical clustering identified shared metabolic states across enzyme knockdowns. Notably, the HPI 80%, HPI 100%, and G6PDH 100% knockdown models shared a common metabolic state under CCM conditions across both representation learning and clustering methods. The reduced number of shared metabolic states in the CCM condition indicates that CRC cells grown in CCM utilize more varied strategies to counter metabolic perturbations, even when those perturbations target the same enzyme or pathway. This finding represents a unique advantage to our workflow: only by parsing the full solution space are we able to identify metabolic solutions where knockdowns of different enzymes – sometimes in different pathways – can lead to similar outcomes. Our application of flux sampling revealed the presence of metabolic diversity both between and within enzyme knockdowns. These diverse samples could be used in studies of tumor heterogeneity, as it is evident that one perturbation can result in a range of metabolic adaptations. This is of particular importance in CRC, as extrinsic and intrinsic factors have been shown to diversify CRC cell metabolism, creating tumor subtypes which can affect the degree of treatment resistance, influence treatment vulnerabilities, and act as clinical biomarkers ^24,25,29^. Understanding the diverse metabolic outcomes of perturbation is therefore an essential tool to understand and treat CRC. In addition to allowing us to draw conclusions about tumor heterogeneity, these findings have biological implications for the metabolic relationship between CAFs and tumor cells.

Identification of differences in the shared metabolic states for CCM condition compared to the CRC condition exemplifies a broader finding: CAF secretions in the CCM play a functional role in determining what metabolic adaptations CRC cells take after perturbation. Summed pathway fluxes revealed elevated glycolytic flux in CCM models, consistent with literature showing that CAFs promote glycolysis in tumor cells^30^. Additionally, changes in flux through the PPP were more pronounced in CCM models, with some knockdowns showing reversed flux direction toward reactants. This metabolic behavior may be partially explained by the more homogeneous metabolic states seen in CRC media. Biologically, this shift could suggest increased production of glycolytic intermediates and a preference for NADPH and ribose-5-phosphate (R5P) generation, which are critical for redox balance and biosynthesis^31^. The magnitude of negative PPP flux in CCM may reflect CAF-induced promotion of R5P synthesis. R5P functions as a metabolic checkpoint between glycolysis and PPP flux, and its modulation may help explain the observed shifts^32^. In some models, reactions in the oxidative branch of the PPP were upregulated, potentially to boost NADPH production and mitigate oxidative stress, which may be accompanied by decreased glycolysis in the same models. Moreover, the GAPDH knockdown model behaved more like ENO, PYK, or PGAM knockdowns in the CCM condition while aligning with HPI, HK, G6PDH, or ALDO knockdowns in CRC condition. This suggests that the presence of CAF secretions in CRC cell media may shift GAPDH-related responses. Future work could further investigate the role of GAPDH in CAF-influenced tumor metabolism.

The biological consequences of these pathway flux shifts within metabolic states were assessed by examining biomass flux. Previous studies have shown that CAFs provide metabolic protection and promote resistance to therapy through metabolic reprogramming^11,33^. Consistent with those observations, we demonstrated that partial knockouts of glycolytic enzymes can result in a similar metabolic state to complete knockdowns and can lead to a complete reduction in biomass flux. This finding is supported by published studies, which indicate that partial inhibition of glycolysis may result in a reduction in tumor cell viability and proliferation^36,36^. Furthermore, prior work emphasizes the importance of including TME components in therapeutic development and drug testing, as we demonstrate that cell-cell interactions can shape metabolic outcomes post perturbation^37,38,39^. Additionally, CAFs are functionally diverse, and multiple CAF subtypes have been shown to contribute to treatment resistance^40,41^. Studies have shown that CAFs can contribute to resistance to chemotherapy, targeted therapies, and immune checkpoint inhibitors in multiple cancer types^42,43^. Consistent with this literature, in approximately half of the knockdown models predicted biomass flux was lower in CRC media compared to CCM, indicating that CAFs buffer against the impact of enzyme inhibition. Collectively, these results suggest that CAFs can provide a protective effect against metabolic perturbation, even in response to the most impactful enzyme knockdowns.

This conclusion could have implications for the development of therapeutics targeting KRAS-mutant CRC. As the KRAS-mutation can make CRC cells more aggressive and more prone to drug resistance, discovering new compounds is essential, and these results could be advantageous in this search^44,45^. It is evident from our findings that CCM can induce metabolic heterogeneity after perturbation. In the context of drug development, this means that the results of drug testing could be altered if the experiments are completed in CCM or normal media. Already, drug discovery investigations are considering the impact of components of the TME on drug response^46,47^. This inclusion is particularly important when we consider that no knockdown model cluster in CCM shared a metabolic state with a knockdown model in CRC in the representation learning projection. While pathway-level analysis revealed that directional changes in pathway flux were sometimes consistent across media conditions, the magnitude of change was often distinct, meaning that while one drug may impact the same pathways whether the CRC cells are in the presence or absence of CAFs, the degree of impact can vary. Particularly in CRC, where glycolysis and glutaminolysis pathways are known to be altered, the conclusion that there is a media-dependent shift could have meaningful consequences^11,48,49^.

We recognize some limitations of this study. The analysis employed a single metabolic model representing KRAS-mutant CRC cells in the presence of CAF-released factors, rather than a multi-compartment model that explicitly simulates cell-cell metabolite exchange. While this simplification enables clearer interpretation of metabolic trends, it limits the ability to analyze transport and exchange reactions. Therefore, we focused on central carbon pathways (glycolysis, glutaminolysis, PPP, and the TCA cycle) where metabolic reprogramming is well characterized. While our goal was to elucidate the metabolic interactions between CAFs and CRC cells, the use of a conditioned media to represent this relationship means we are limited in the conclusions we can draw about CAF-tumor crosstalk. This bi-directional crosstalk is well described in literature, where CAFs undergo their own metabolic changes in addition to altering the metabolic processes of CRC cells^50,51^. We can reasonably conclude that our CAF-conditioned media contains secretions from fibroblasts, which interact with the CRC cells. This means that our model represents, unidirectionally, the impact of CAF-released factors on CRC cells. Conclusions from this model are therefore most meaningful in the context of CRC cell metabolic alterations both in expanding our understanding of how CRC cells respond to perturbation in the context of the TME, and in the development of metabolism-targeting therapeutics. However, further research is necessary to understand what mechanisms drive the observed shifts toward specific metabolic states, as these shifts could be governed by metabolic need, availability of CAF-derived resources, or other regulatory factors. Future works could expand these conclusions to bi-directional CAF-tumor crosstalk by constraining a similar model with results from co-culture experiments, or by designing a multi-compartment model.

Our decision to use a focused model structure also means that the model did not include pathways outside of central carbon metabolism such as lipid metabolism, which is known to be altered in CRC and influenced by CAF activity^52^. Future work could employ genome-scale metabolic modeling to capture broader pathway effects and incorporate multiple cell types and interaction mechanisms. Finally, our results support the idea that targeting metabolic crosstalk or disrupting CAF-induced metabolic adaptations could provide therapeutic benefit; however, additional work needs to be done to determine if the identified metabolic state shifts are therapeutically targetable.

## Conclusion

Our study underscores the critical role of CAFs in shaping metabolic response to perturbation in KRAS-mutant CRC cells. The results emphasize the importance of including TME components like CAFs when evaluating potential metabolic therapies. Incorporating such biological complexity into computational models could significantly improve the predictive power of therapeutic strategies. More broadly, this systems-level approach offers a promising framework for drug discovery and development, particularly in the context of metabolic vulnerabilities shaped by heterogeneity and tumor-stroma interactions.

## Methods

### Model construction and constraint-based modeling

We utilized a previously published, experimentally informed model of central carbon metabolism, as described by Wang et al^14^. The model consists of 74 reactions, including one reaction to represent cellular biomass production. Models were experimentally informed to replicate two growth conditions: KRAS-mutant CRC cells grown in matched, colorectal cancer media, and KRAS-mutant CRC cells grown in CAF-conditioned media. For the CAF-conditioned media condition, CAFs were cultured and then removed from media before introducing CRC cells. The cell growth (biomass reaction flux), glucose and glutamine uptake, and lactate secretion rates were initially set based on experimental data and were used to constrain the models, as described in Wang et al. The biomass reaction is used as a proxy for growth the objective function in flux balance analysis, which was performed to identify the initial enzyme knockdowns of interest. We then perturbed the base model to simulate enzyme knockdowns. For 100% knockdowns, the upper and lower bounds for the reaction mediated by the enzyme being perturbed were constrained to 0 mM/h. For partial enzyme knockdowns, the bounds were changed relative to the unperturbed model. For example, for HK 80%, the upper bounds of the reaction mediated by HK were limited to 20% of the flux predicted in the unperturbed model. After simulating the enzyme knockdowns, flux sampling was applied to the models.

### Flux sampling

To perform flux sampling, maximizing flux through the biomass reaction was removed as the objective function. This allowed the reaction fluxes to vary outside of the range limited by optimizing biomass, producing the full range of possible values for each reaction. Artificially-centered hit-and-run (ACHR) sampling was used as the sampling method. ACHR sampling first estimates the center of the solution space, which is then revised as samples are generated^53,54^. The new center estimate then informs the next sample, so the solution space is covered in fewer steps compared to traditional hit-and-run sampling^55,56^. A total of 1000 samples were generated for each reaction in each model. The number of samples was based on the small size of the model, the narrow deviation around the mean from the biomass sampling distribution **(Supplemental Table 2)**, and past works^57^. Both constraint-based modeling and flux sampling were performed in MATLAB R2024a using the COBRA Toolbox v3.0.

### Dimensionality reduction

Representation learning was performed in Python (v3.11.6) and was applied to the flux sampling output following the approach described by Tavakoli et al^15^. Representation learning utilizes neural networks to transform multi-dimensional inputs into lower dimension features^28^. Our representation learning approach utilized Siamese neural networks, which interpret input matrices as images and weigh each datapoint in the context of the full matrix to identify features^58^. Siamese neural networks (SNNs) are chosen for their ability to learn the encoded representations of input vectors and output projected representations with meaningful and quantifiable differences^59,60^. The approach therefore outputs a low-dimensional representation of the model data that uses an optimization function to minimize the distance between projections and captures the similarities and differences between the inputs. The distance between these representations of the models can be calculated to quantify the similarity between them.

Unique to this investigation, the input matrices were 1000 rows x 74 columns, representing each of the 1000 samples for each of the 74 metabolic reactions. We produced this matrix under each of the two conditions (KRAS-Mut CCM and KRAS-Mut CRC). Therefore, each sample from each knockdown and condition is mapped into 2-D space, allowing the SNN to learn from the relationships between each flux sample for each enzyme knockdown. SNNs are trained on the entire dataset using triplet loss, which requires explicit labelling to minimize the distance between an anchor data point and a point from the same group, while maximizing the difference between the anchor and a point from a different group^61^. Data are organized and labelled based on their enzyme knockdown and media condition. We employ this form of supervised learning to capture the features of the high-dimensional dataset, while still considering the known labels that separate the data.

Dimensionality reduction is particularly useful in this context, where otherwise quantifying the similarities and differences between 1000 sampling vectors for 26 different labelled groups would be infeasible. Representation learning is advantageous because it projects the original input into a lower-dimensional space, rather than outputting a new rendition of the input data. This means that conclusions we draw about these projections can be applied to the flux sampling vectors themselves. Similar techniques have been successfully used with single-cell cytometry data to characterize cellular composition, retaining the local and global relationships between cell types^62^.

### Statistical Analysis

Statistical analysis was performed in R (v4.4.1). We calculate the centroid of the sample points for each knockdown within each condition as the mean coordinate values from their mapped projection. Similarities between knockdown model clusters are therefore defined by the overlap between clusters and the Euclidean distance between their centroids. Variance within a model cluster was defined as the average Euclidean distance between the cluster centroid and each sampling point.

Percent overlap between projected model clusters was used to define a shared metabolic state. Percent overlap was determined by calculating the convex hull (the smallest shape that contains a set of points) for each model cluster. Then, the area overlapping between two convex hulls was measured. This overlapping area was subtracted from the total area of the convex hulls. The overlapping area was divided by the total area, minus the overlapping area, and multiplied by 100% to obtain the percent overlap. Percent overlap calculations were performed using the *geometry* package (v0.5.2).

Dendrogram clustering was performed based on Euclidean distance of the mean flux values for each reaction in each knockdown model and describes the hierarchical relationship between model flux predictions. To determine the hierarchical relationship, complete-linkage (farthest neighbor) clustering is used, which is an agglomeration method that calculates the furthest distance between the Euclidean distance matrices in order to emphasize the differences between predicted model fluxes. Media condition heatmaps and clustering were performed using the *ComplexHeatmap* package (v2.22.0). For the sum pathway flux, reactions were grouped by pathway for each sample in each model. The mean and standard deviation of all samples within a model were calculated. Biomass flux distributions were compared using the Kolmogorov-Smirnoff test, implemented with the *stats* package (v4.4.1).

## Data Availability

All relevant datasets and scripts for analysis are available on GitHub at: https://github.com/FinleyLabUSC/CRC_Flux_Sampling

## Author Contributions

E.E. conducted all metabolic modeling and computational analyses. N.T. and H.C. contributed to scientific discussions of results. E.E. and S.D.F. contributed to manuscript writing and scientific discussions of results. S.D.F. was responsible for the scientific direction of this work. All authors reviewed and provided feedback on the manuscript.

## Acknowledgements

We would like to thank the members of the Finley Lab at USC for the deep scientific discussions and critical feedback. We acknowledge funding support from the USC Center for Computational Modeling of Cancer.

