## Supplemental Information for "Cancer-associated fibroblasts drive metabolic heterogeneity in KRAS-mutant colorectal cancer cells"

**Supplemental Figure 1. Distributions of predicted flux values for reactions in glutaminolysis, glycolysis, the pentose phosphate pathway, and the tricarboxylic acid cycle.** Sampled flux (mM/h) through the reaction for each model under the CCM condition (left, filled) and the CRC condition (right, open). Some reaction distributions are single values (represented by dash) while others comprise wider distributions (boxplots).

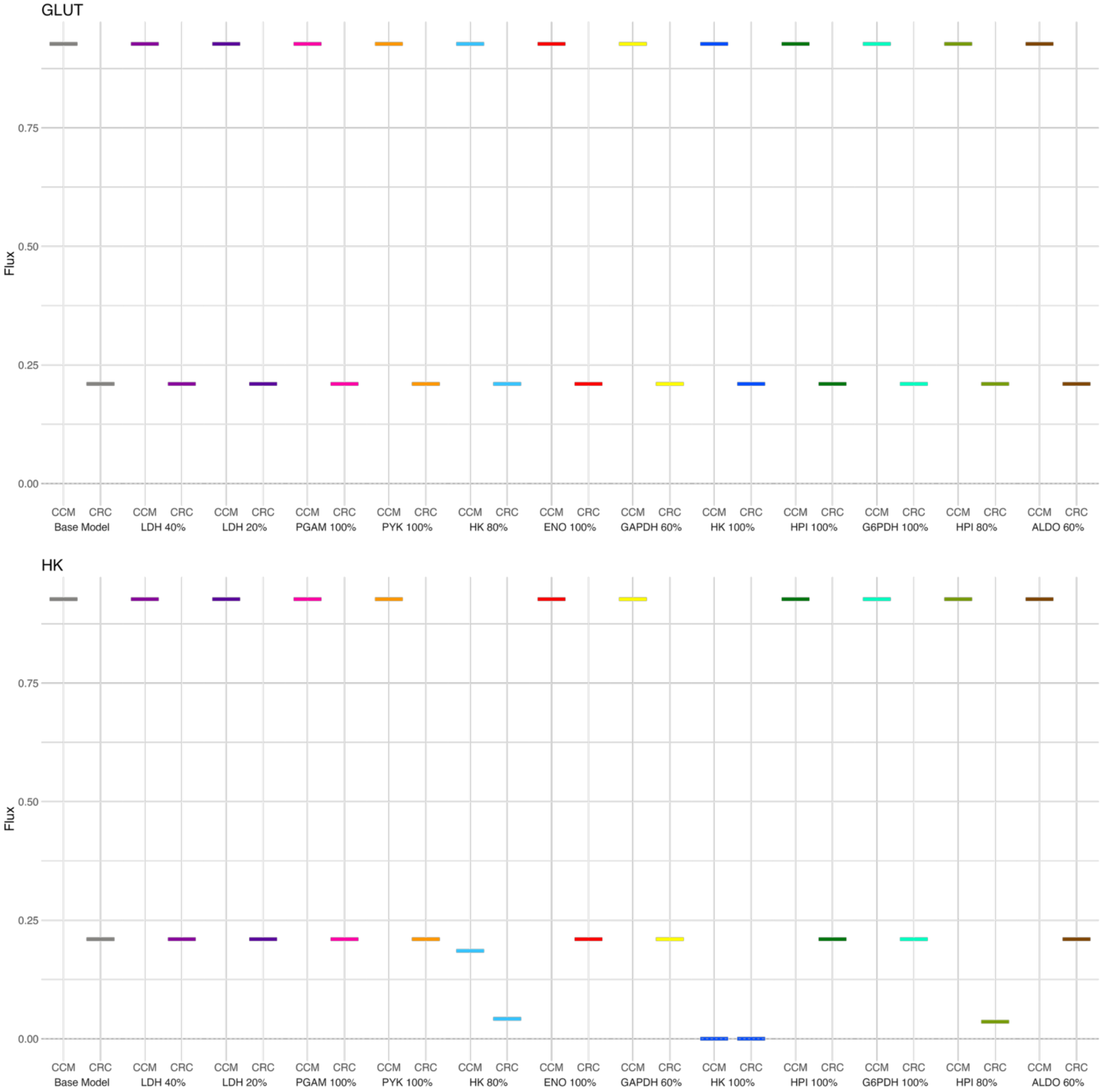

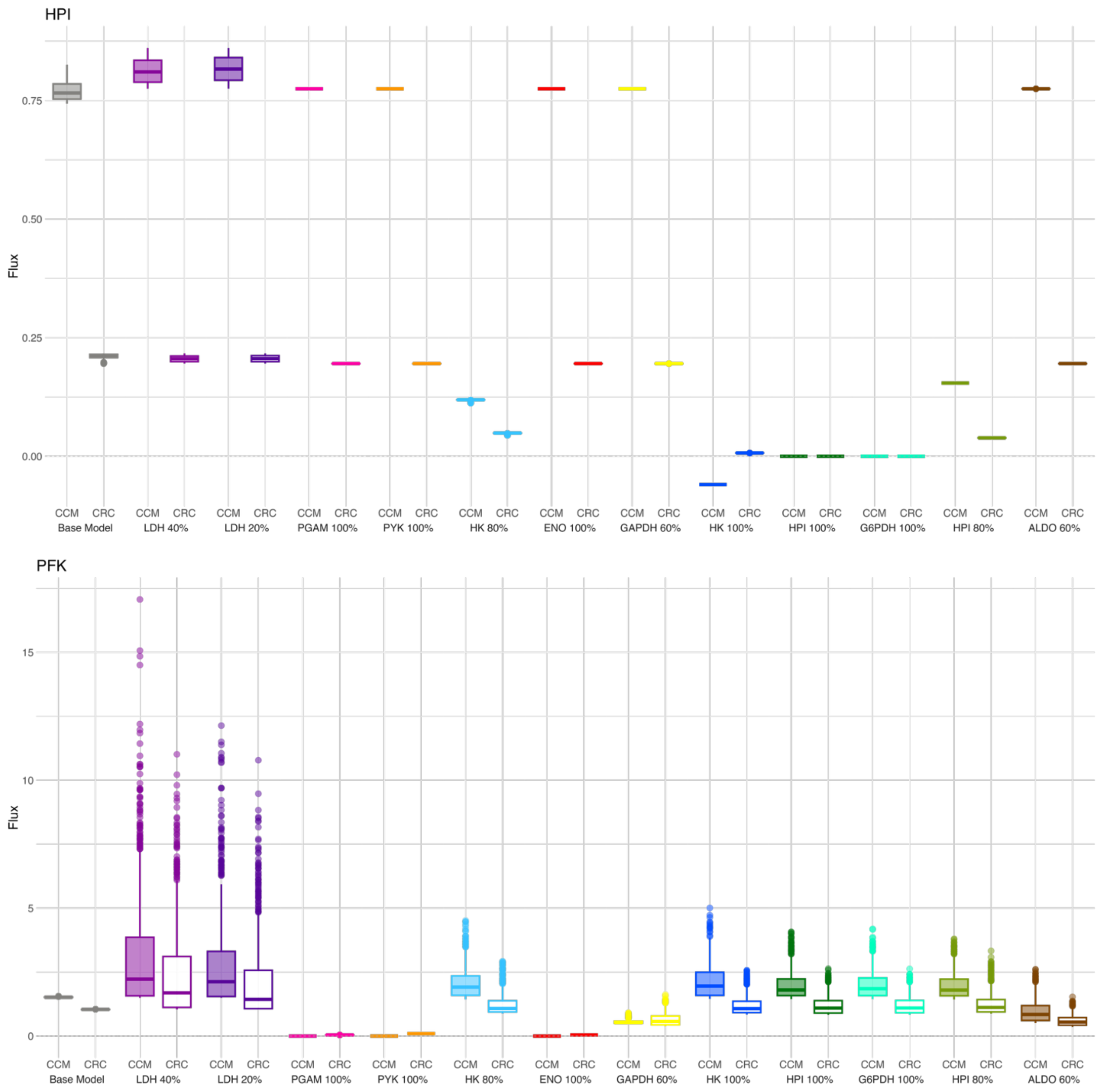
HPI
Flux
0.75
0.50
0.25
0.00
CCM CRC
Base Model
LDH 40%
LDH 20%
PGAM 100%
PYK 100%
HK 80%
ENO 100%
GAPDH 60%
HK 100%
HPI 100%
G6PDH 100%
HPI 80%
ALDO 60%
PFK
15
10
5
0

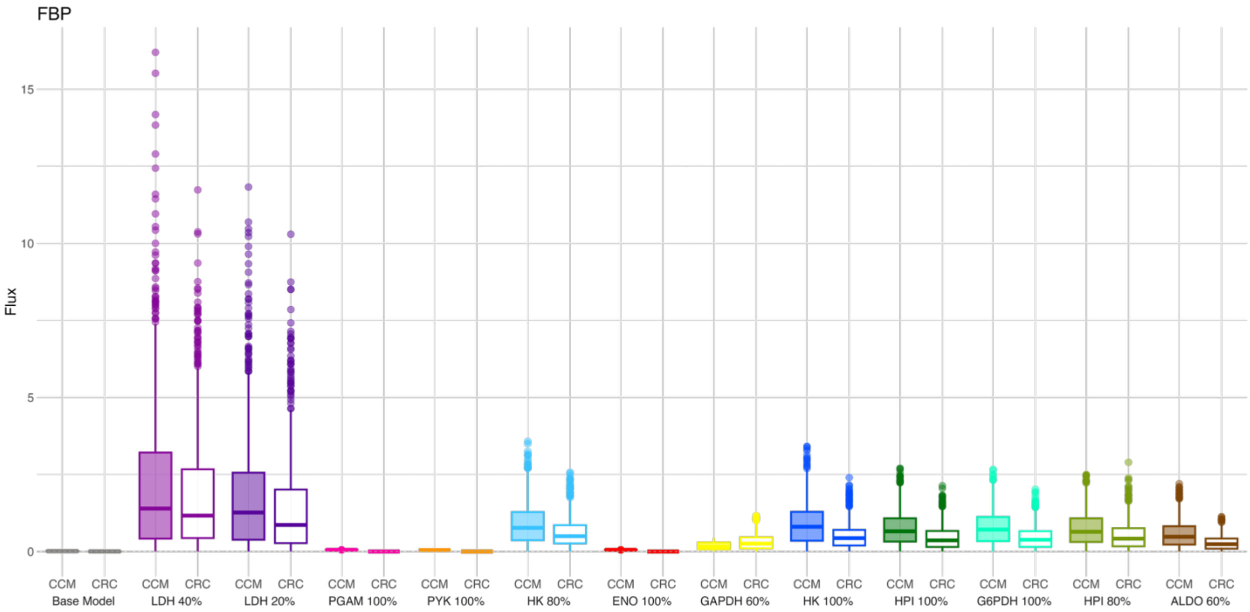

FBP
Flux
15
10
5
0
CCM CRC Base Model
CCM CRC LDH 40%
CCM CRC LDH 20%
CCM CRC PGAM 100%
CCM CRC PYK 100%
CCM CRC HK 80%
CCM CRC ENO 100%
CCM CRC GAPDH 60%
CCM CRC HK 100%
CCM CRC HPI 100%
CCM CRC G6PDH 100%
CCM CRC HPI 80%
CCM CRC ALDO 60%


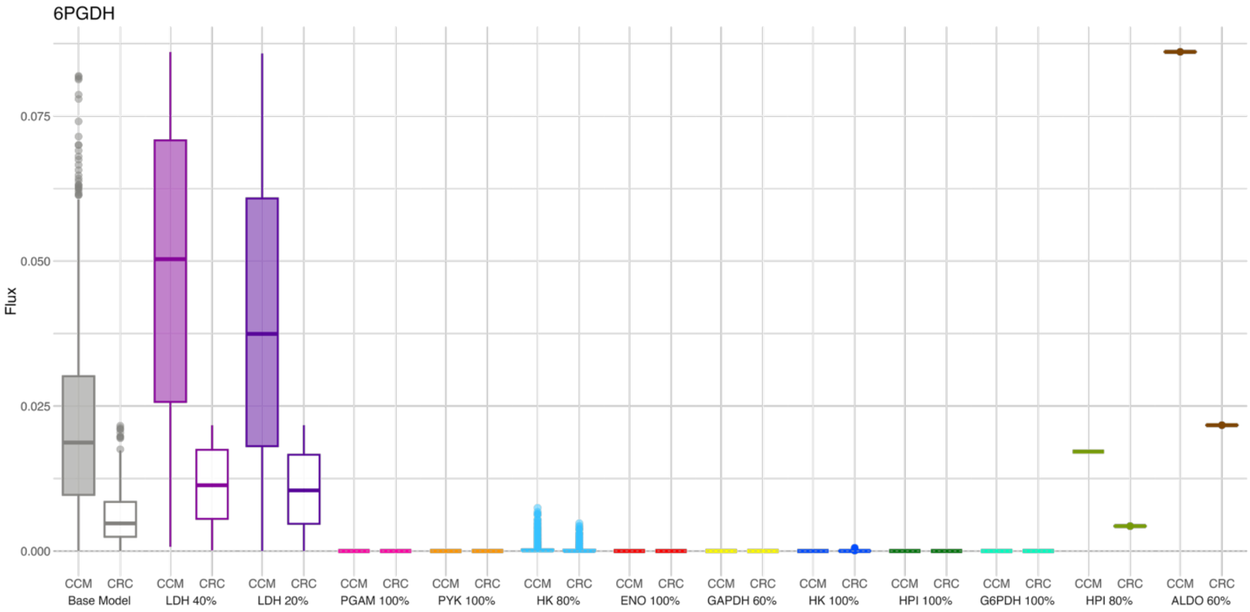

6PGDH
Flux
0.075
0.050
0.025
0.000
CCM CRC Base Model
CCM CRC LDH 40%
CCM CRC LDH 20%
CCM CRC PGAM 100%
CCM CRC PYK 100%
CCM CRC HK 80%
CCM CRC ENO 100%
CCM CRC GAPDH 60%
CCM CRC HK 100%
CCM CRC HPI 100%
CCM CRC G6PDH 100%
CCM CRC HPI 80%
CCM CRC ALDO 60%

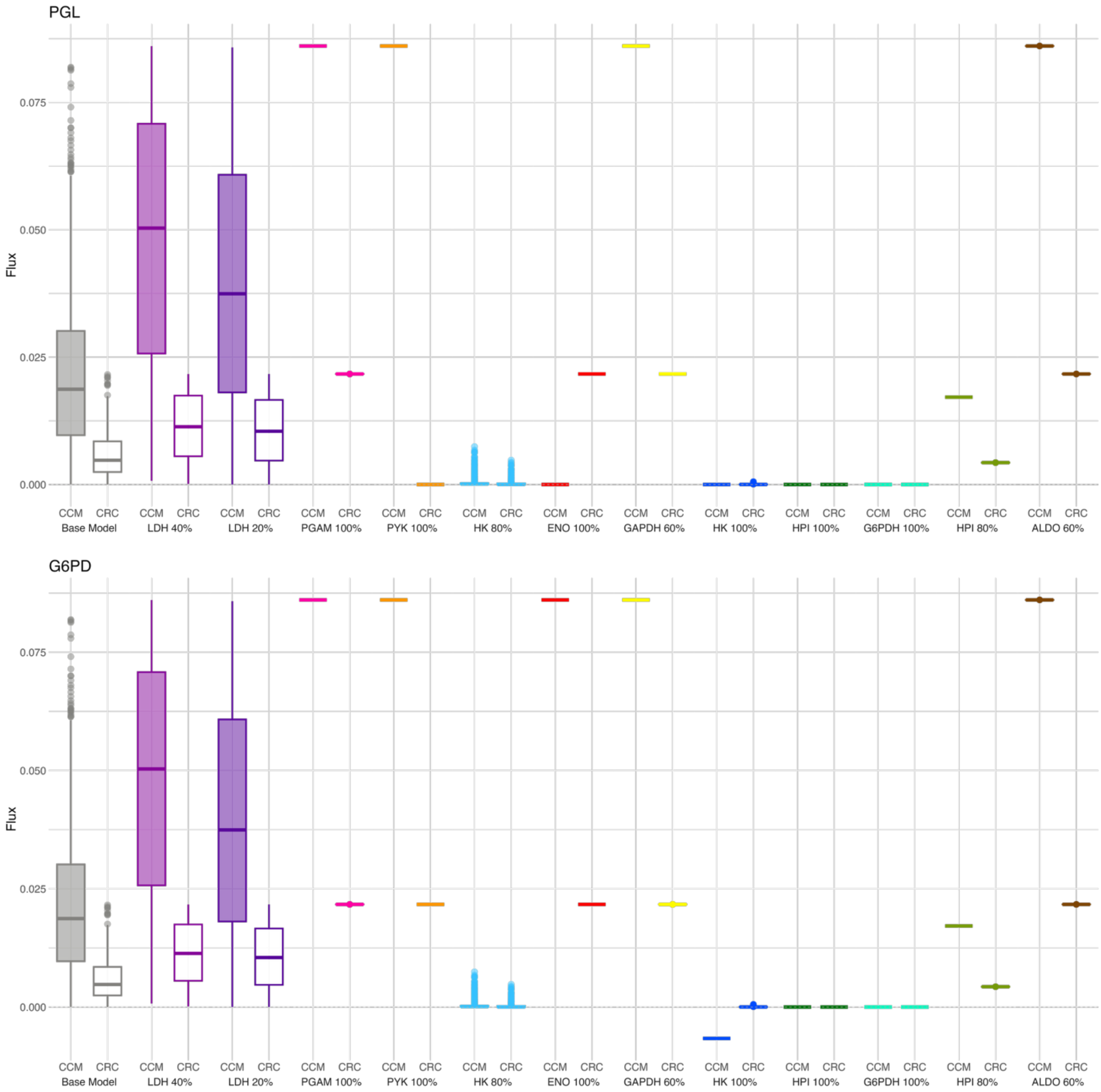
PGL
Flux
0.000
0.025
0.050
0.075
CCM CRC
Base Model
LDH 40%
LDH 20%
PGAM 100%
PYK 100%
HK 80%
ENO 100%
GAPDH 60%
HK 100%
HPI 100%
G6PDH 100%
HPI 80%
ALDO 60%
G6PD
Flux
0.000
0.025
0.050
0.075
CCM CRC
Base Model
LDH 40%
LDH 20%
PGAM 100%
PYK 100%
HK 80%
ENO 100%
GAPDH 60%
HK 100%
HPI 100%
G6PDH 100%
HPI 80%
ALDO 60%

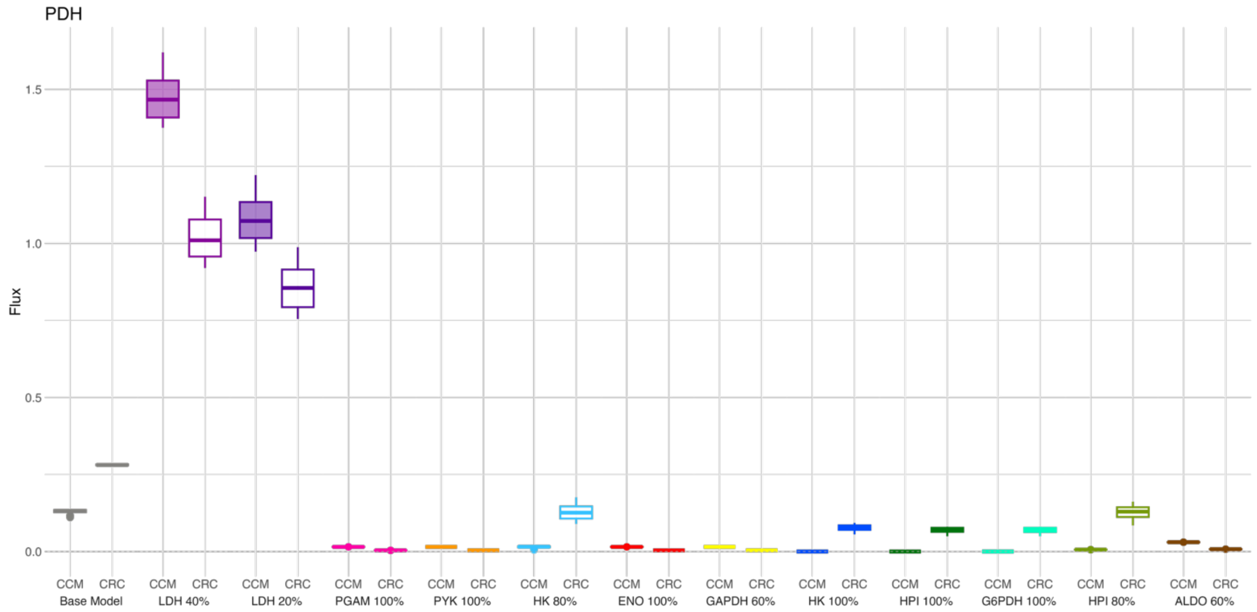
PDH
Flux
1.5
1.0
0.5
0.0
CCM CRC
Base Model
LDH 40%
LDH 20%
PGAM 100%
PYK 100%
HK 80%
ENO 100%
GAPDH 60%
HK 100%
HPI 100%
G6PDH 100%
HPI 80%
ALDO 60%

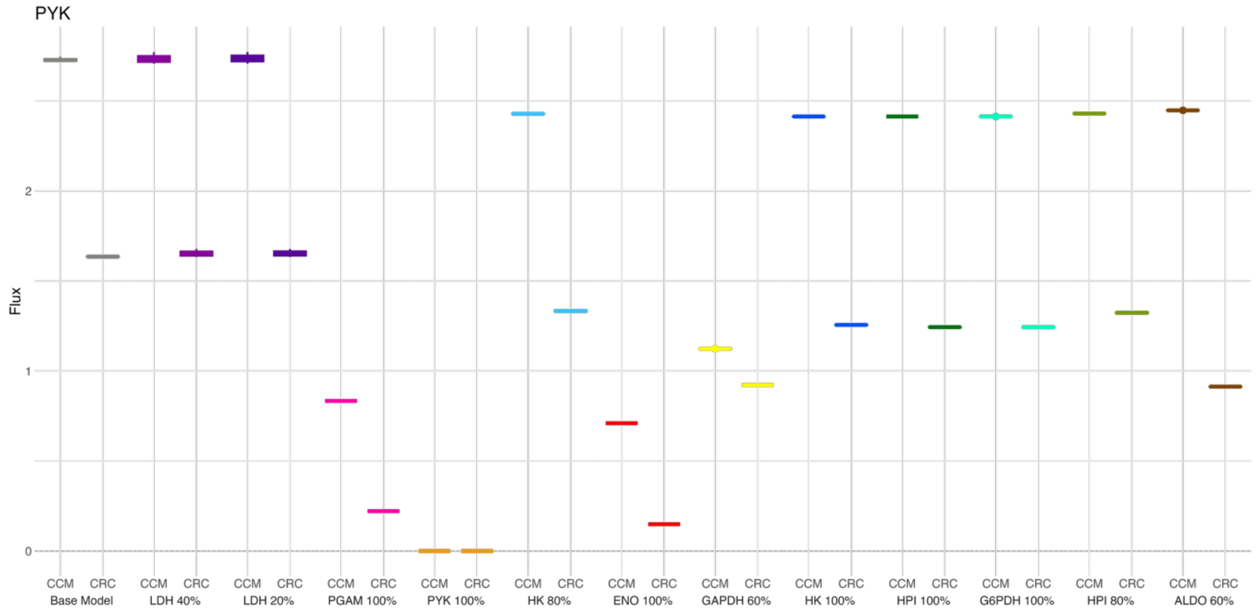
PYK
Flux
2
1
0
CCM CRC
Base Model
LDH 40%
LDH 20%
PGAM 100%
PYK 100%
HK 80%
ENO 100%
GAPDH 60%
HK 100%
HPI 100%
G6PDH 100%
HPI 80%
ALDO 60%

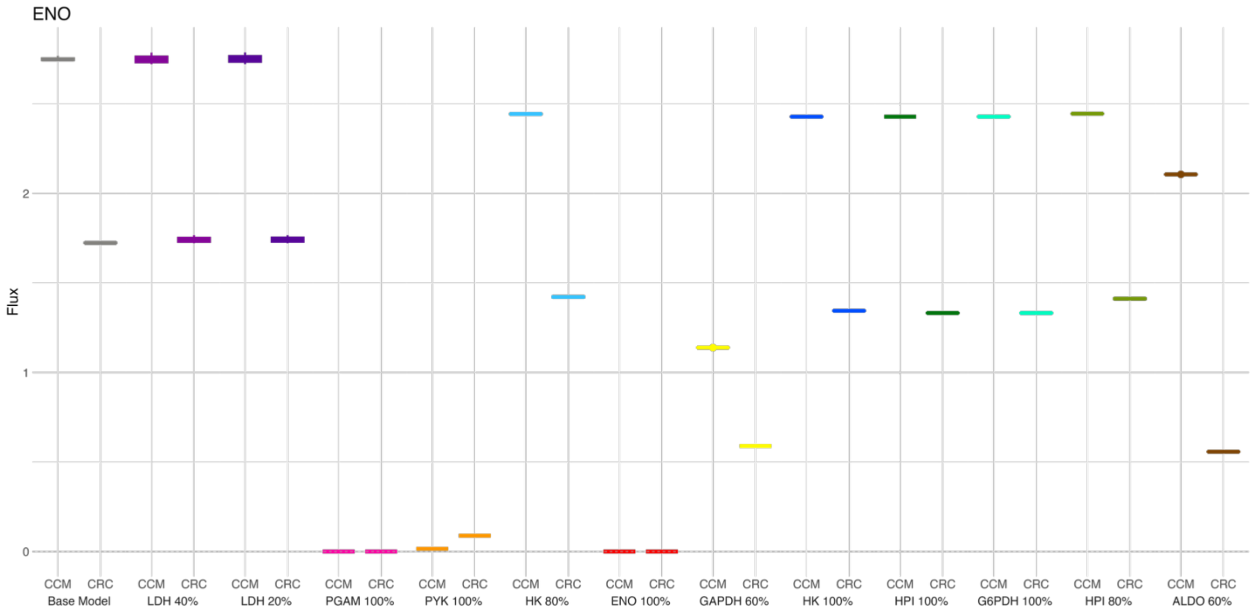
ENO
Flux
2
1
0
CCM CRC
Base Model
CCM CRC
LDH 40%
CCM CRC
LDH 20%
CCM CRC
PGAM 100%
CCM CRC
PYK 100%
CCM CRC
HK 80%
CCM CRC
ENO 100%
CCM CRC
GAPDH 60%
CCM CRC
HK 100%
CCM CRC
HPI 100%
CCM CRC
G6PDH 100%
CCM CRC
HPI 80%
CCM CRC
ALDO 60%

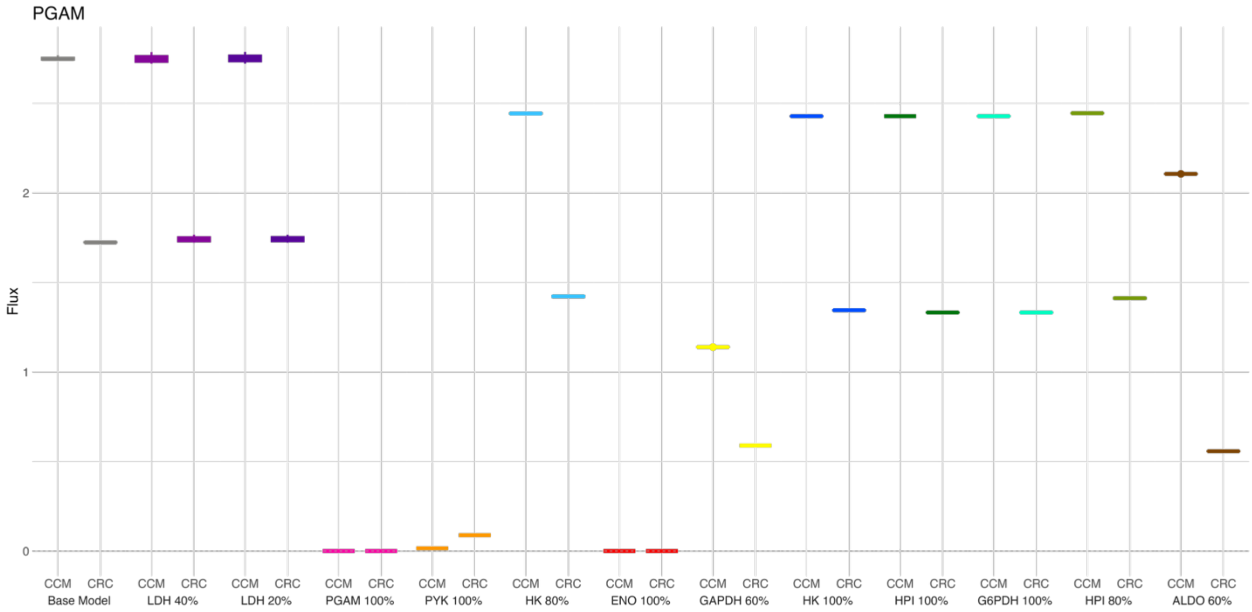
PGAM
Flux
2
1
0
CCM CRC
Base Model
CCM CRC
LDH 40%
CCM CRC
LDH 20%
CCM CRC
PGAM 100%
CCM CRC
PYK 100%
CCM CRC
HK 80%
CCM CRC
ENO 100%
CCM CRC
GAPDH 60%
CCM CRC
HK 100%
CCM CRC
HPI 100%
CCM CRC
G6PDH 100%
CCM CRC
HPI 80%
CCM CRC
ALDO 60%

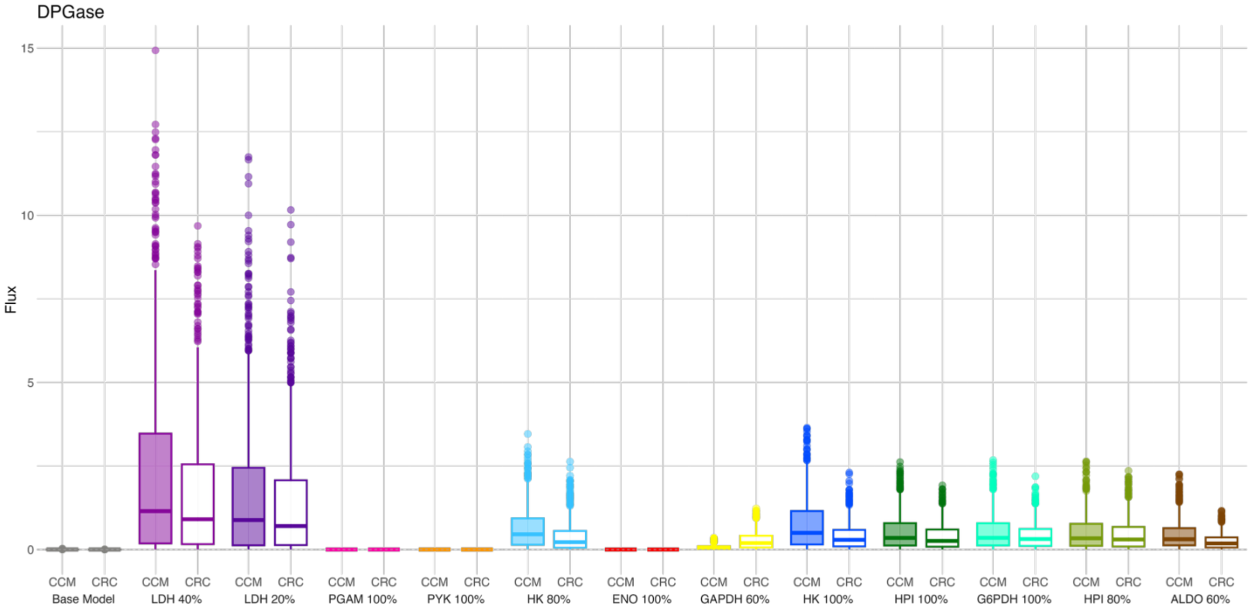
DPGase
Flux
15
10
5
0
CCM CRC
Base Model
CCM CRC
LDH 40%
CCM CRC
LDH 20%
CCM CRC
PGAM 100%
CCM CRC
PYK 100%
CCM CRC
HK 80%
CCM CRC
ENO 100%
CCM CRC
GAPDH 60%
CCM CRC
HK 100%
CCM CRC
HPI 100%
CCM CRC
G6PDH 100%
CCM CRC
HPI 80%
CCM CRC
ALDO 60%

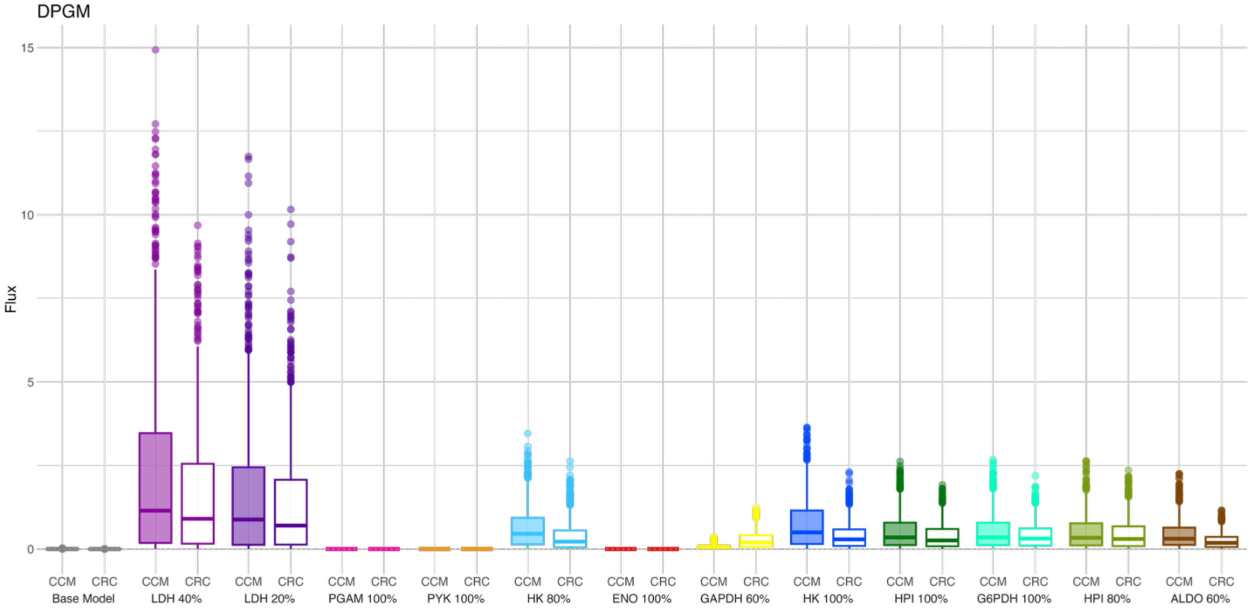
DPGM
Flux
15
10
5
0
CCM CRC
Base Model
CCM CRC
LDH 40%
CCM CRC
LDH 20%
CCM CRC
PGAM 100%
CCM CRC
PYK 100%
CCM CRC
HK 80%
CCM CRC
ENO 100%
CCM CRC
GAPDH 60%
CCM CRC
HK 100%
CCM CRC
HPI 100%
CCM CRC
G6PDH 100%
CCM CRC
HPI 80%
CCM CRC
ALDO 60%

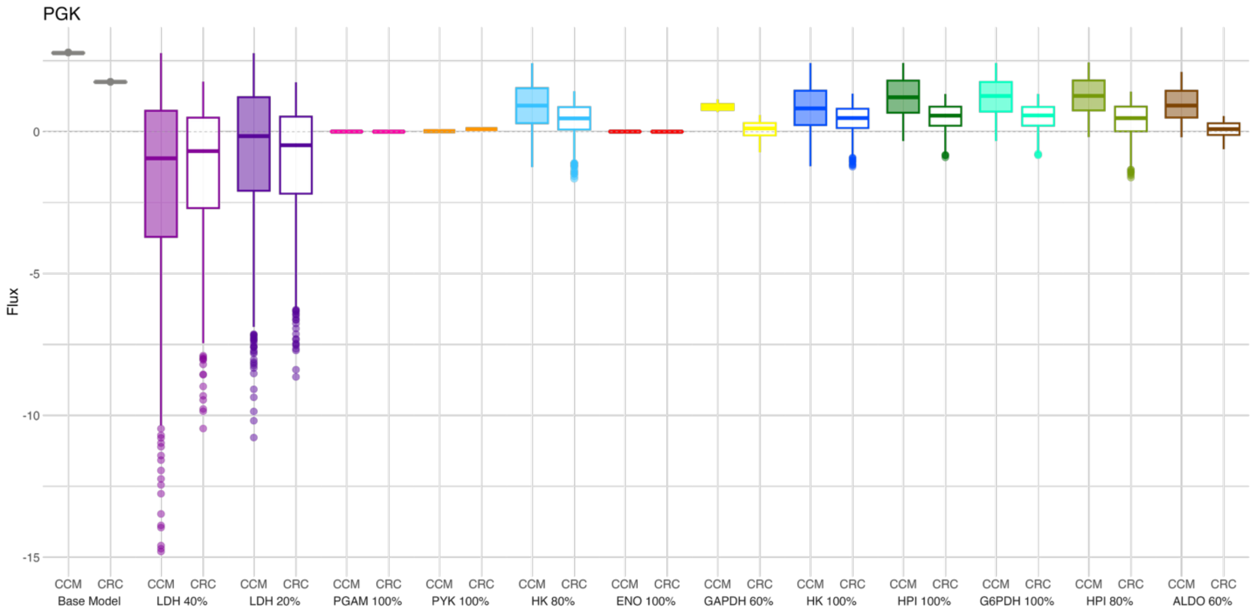
PGK
Flux
0
-5
-10
-15
CCM CRC Base Model
CCM CRC LDH 40%
CCM CRC LDH 20%
CCM CRC PGAM 100%
CCM CRC PYK 100%
CCM CRC HK 80%
CCM CRC ENO 100%
CCM CRC GAPDH 60%
CCM CRC HK 100%
CCM CRC HPI 100%
CCM CRC G6PDH 100%
CCM CRC HPI 80%
CCM CRC ALDO 60%

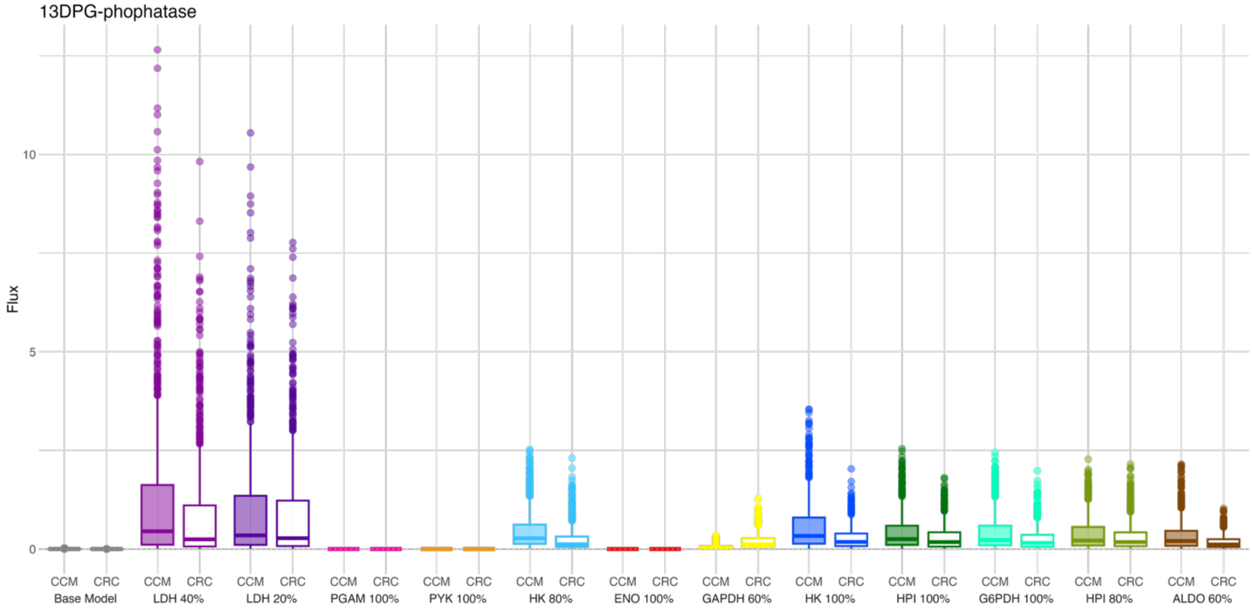
13DPG-phophatase
Flux
10
5
0
CCM CRC Base Model
CCM CRC LDH 40%
CCM CRC LDH 20%
CCM CRC PGAM 100%
CCM CRC PYK 100%
CCM CRC HK 80%
CCM CRC ENO 100%
CCM CRC GAPDH 60%
CCM CRC HK 100%
CCM CRC HPI 100%
CCM CRC G6PDH 100%
CCM CRC HPI 80%
CCM CRC ALDO 60%

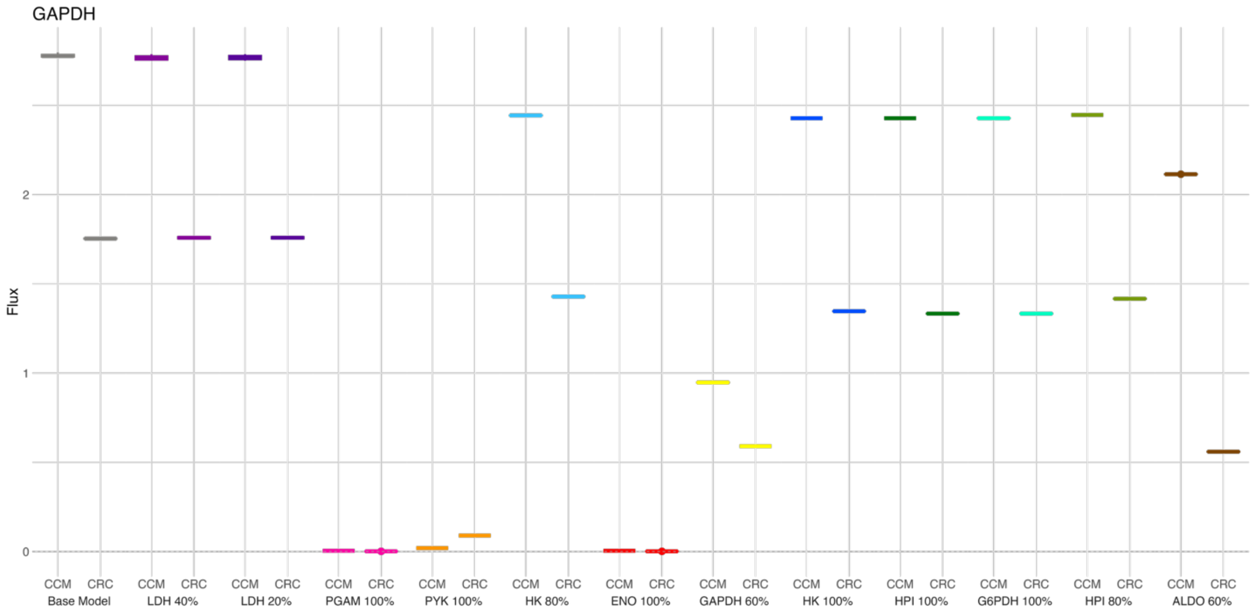
GAPDH
Flux
0
1
2
CCM CRC CCM CRC CCM CRC CCM CRC CCM CRC CCM CRC CCM CRC CCM CRC CCM CRC CCM CRC CCM CRC CCM CRC CCM CRC
Base Model
LDH 40%
LDH 20%
PGAM 100%
PYK 100%
HK 80%
ENO 100%
GAPDH 60%
HK 100%
HPI 100%
G6PDH 100%
HPI 80%
ALDO 60%

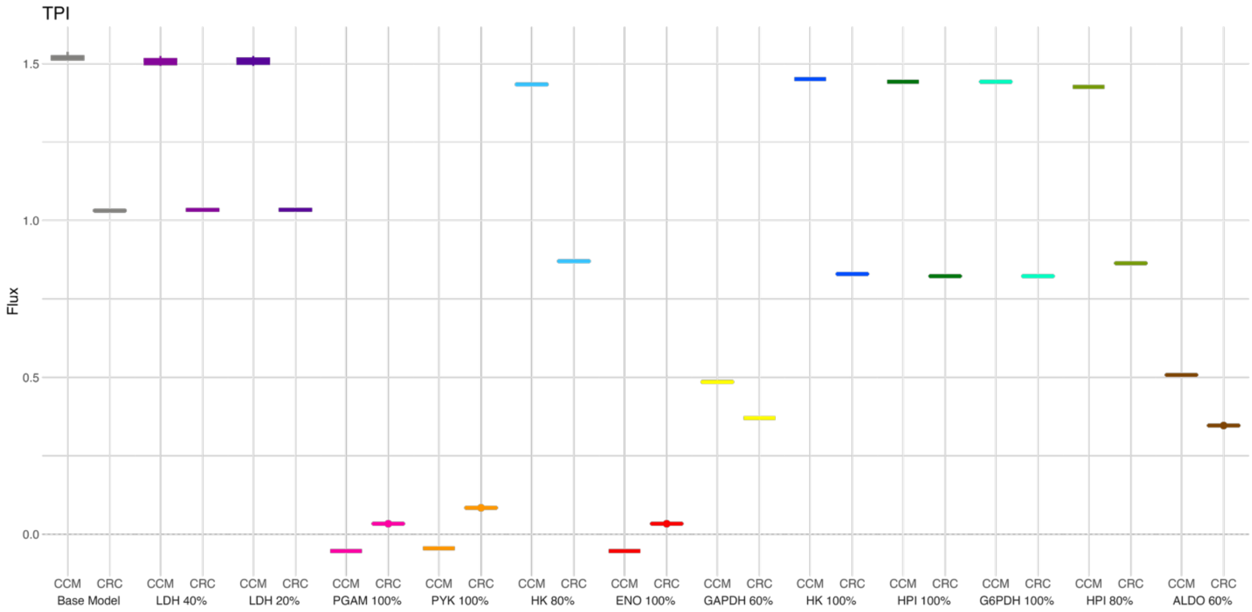
TPI
Flux
0.0
0.5
1.0
1.5
CCM CRC CCM CRC CCM CRC CCM CRC CCM CRC CCM CRC CCM CRC CCM CRC CCM CRC CCM CRC CCM CRC CCM CRC CCM CRC
Base Model
LDH 40%
LDH 20%
PGAM 100%
PYK 100%
HK 80%
ENO 100%
GAPDH 60%
HK 100%
HPI 100%
G6PDH 100%
HPI 80%
ALDO 60%

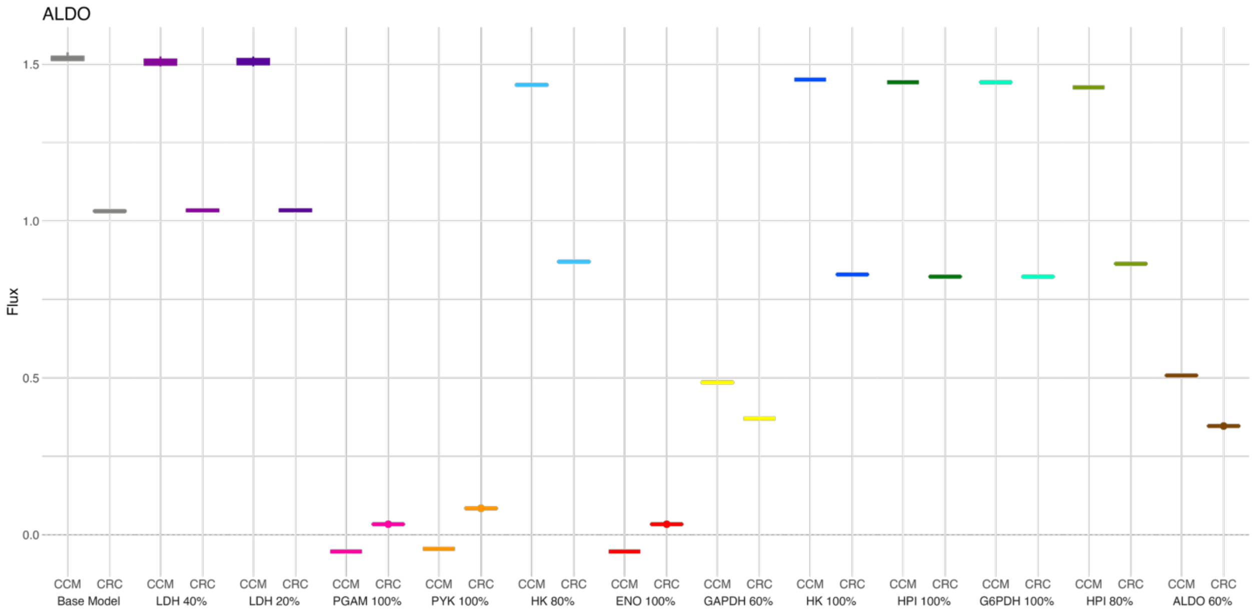
ALDO
Flux
1.5
1.0
0.5
0.0
CCM CRC
Base Model
LDH 40%
LDH 20%
PGAM 100%
PYK 100%
HK 80%
ENO 100%
GAPDH 60%
HK 100%
HPI 100%
G6PDH 100%
HPI 80%
ALDO 60%

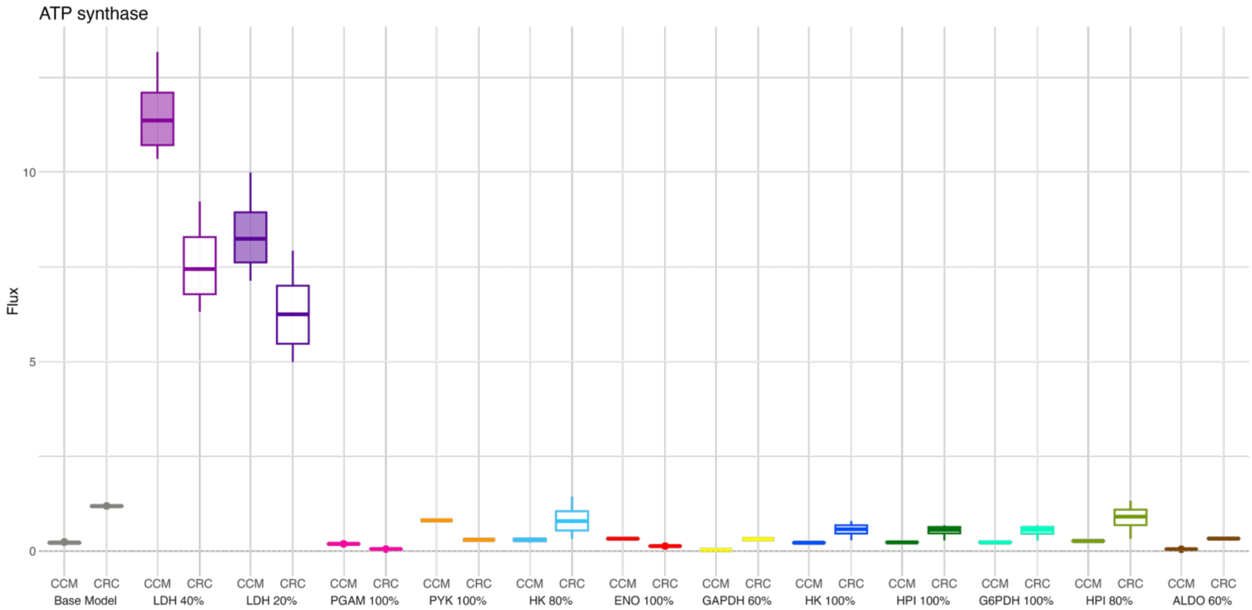
ATP synthase
Flux
10
5
0
CCM CRC
Base Model
LDH 40%
LDH 20%
PGAM 100%
PYK 100%
HK 80%
ENO 100%
GAPDH 60%
HK 100%
HPI 100%
G6PDH 100%
HPI 80%
ALDO 60%

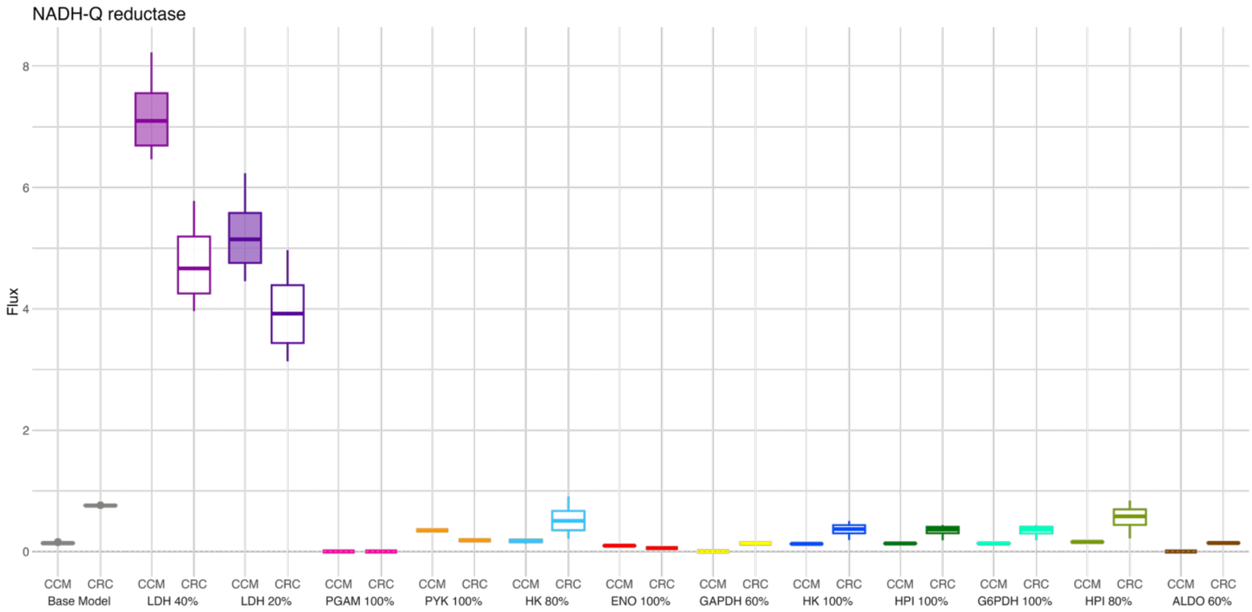
NADH-Q reductase
Flux
0
2
4
6
8
CCM CRC
Base Model
LDH 40%
LDH 20%
PGAM 100%
PYK 100%
HK 80%
ENO 100%
GAPDH 60%
HK 100%
HPI 100%
G6PDH 100%
HPI 80%
ALDO 60%

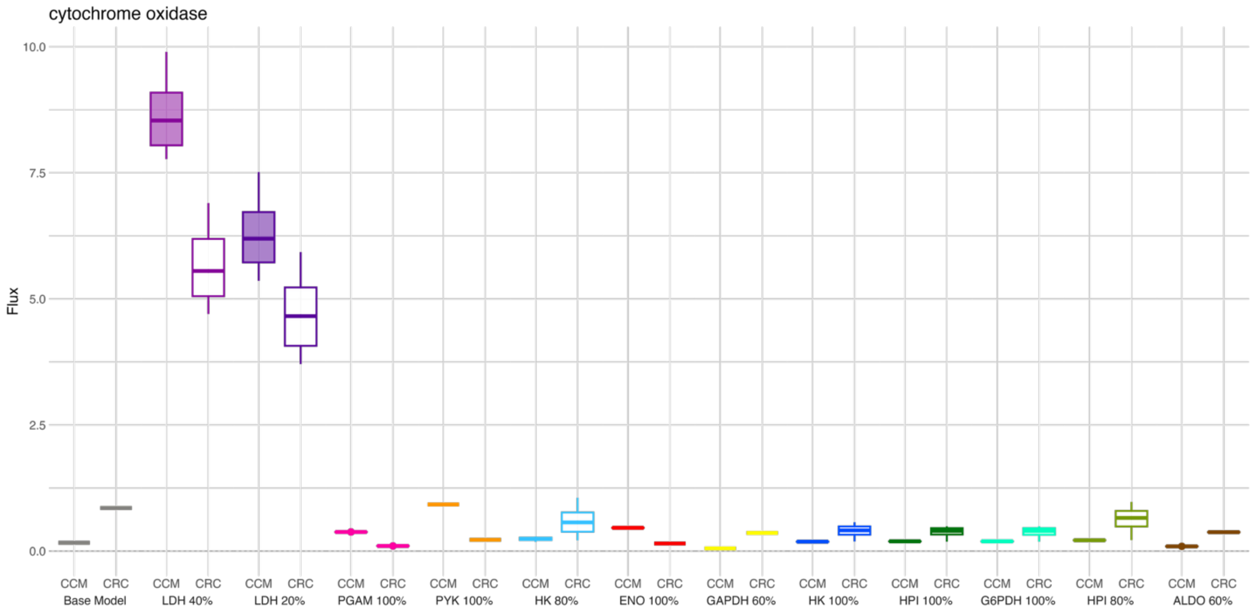
cytochrome oxidase
Flux
0.0
2.5
5.0
7.5
10.0
CCM CRC
Base Model
LDH 40%
LDH 20%
PGAM 100%
PYK 100%
HK 80%
ENO 100%
GAPDH 60%
HK 100%
HPI 100%
G6PDH 100%
HPI 80%
ALDO 60%

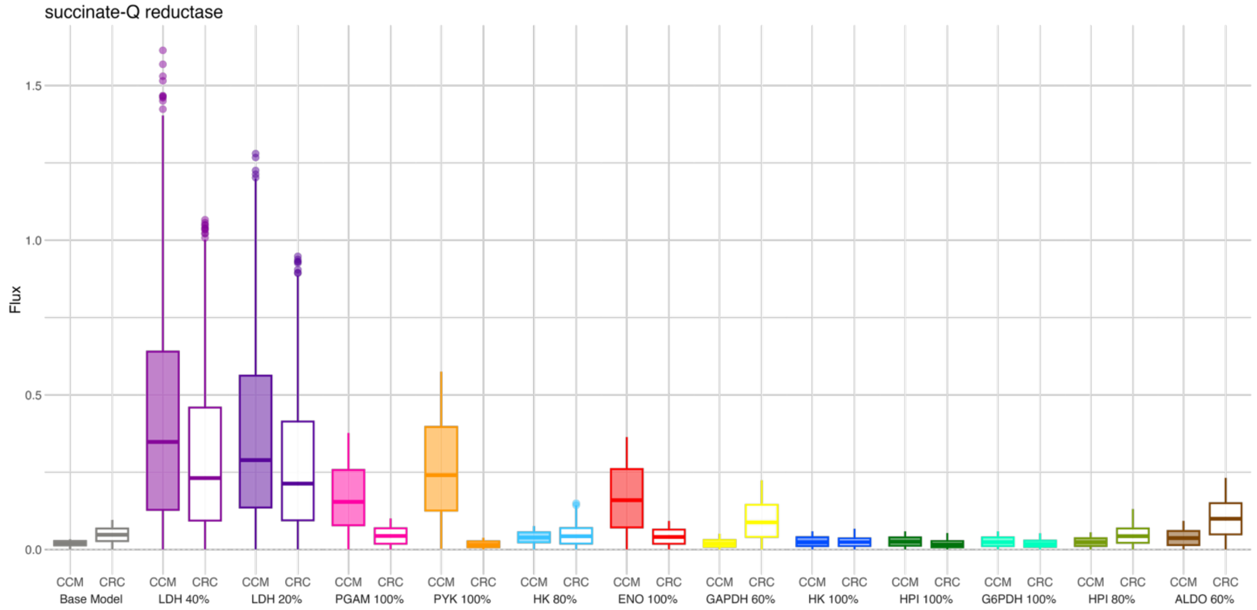
succinate-Q reductase
Flux
1.5
1.0
0.5
0.0
CCM CRC CCM CRC CCM CRC CCM CRC CCM CRC CCM CRC CCM CRC CCM CRC CCM CRC CCM CRC CCM CRC CCM CRC CCM CRC
Base Model LDH 40% LDH 20% PGAM 100% PYK 100% HK 80% ENO 100% GAPDH 60% HK 100% HPI 100% G6PDH 100% HPI 80% ALDO 60%

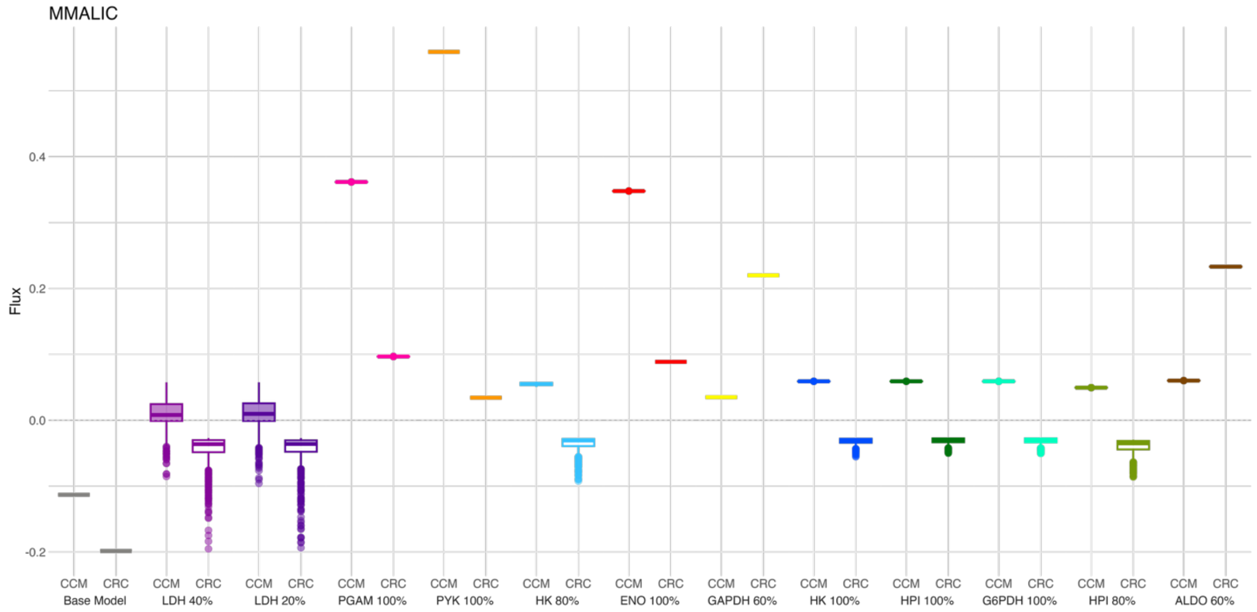
MMALIC
Flux
0.4
0.2
0.0
-0.2
CCM CRC CCM CRC CCM CRC CCM CRC CCM CRC CCM CRC CCM CRC CCM CRC CCM CRC CCM CRC CCM CRC CCM CRC CCM CRC
Base Model LDH 40% LDH 20% PGAM 100% PYK 100% HK 80% ENO 100% GAPDH 60% HK 100% HPI 100% G6PDH 100% HPI 80% ALDO 60%

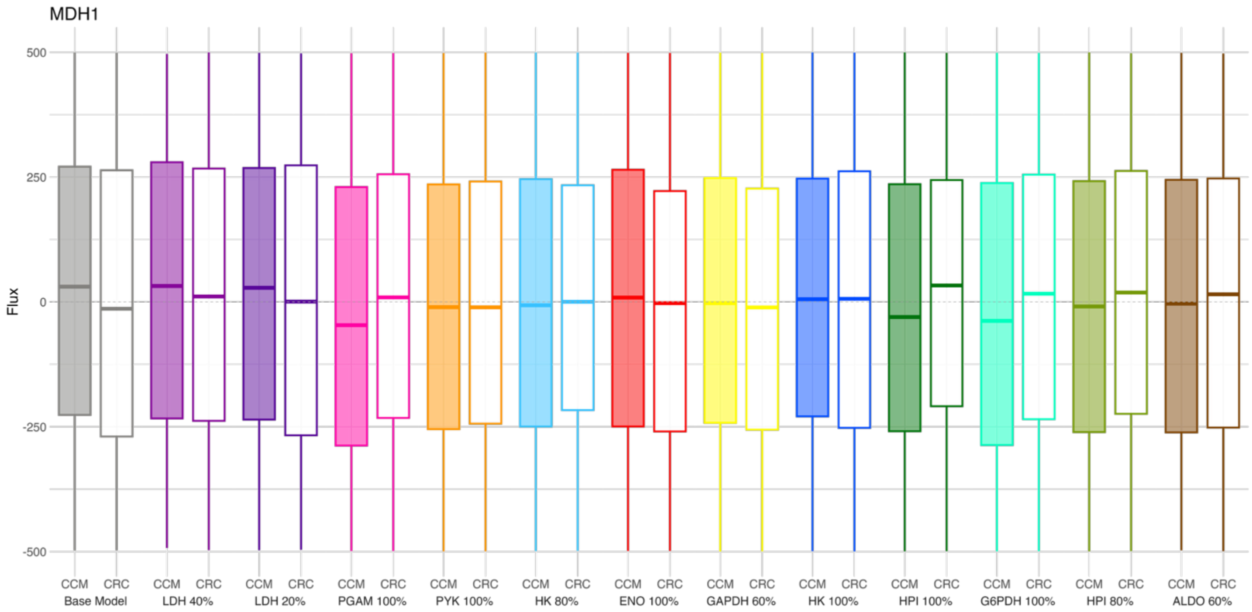
MDH1
Flux
500
250
0
-250
-500
CCM CRC
Base Model
LDH 40%
LDH 20%
PGAM 100%
PYK 100%
HK 80%
ENO 100%
GAPDH 60%
HK 100%
HPI 100%
G6PDH 100%
HPI 80%
ALDO 60%

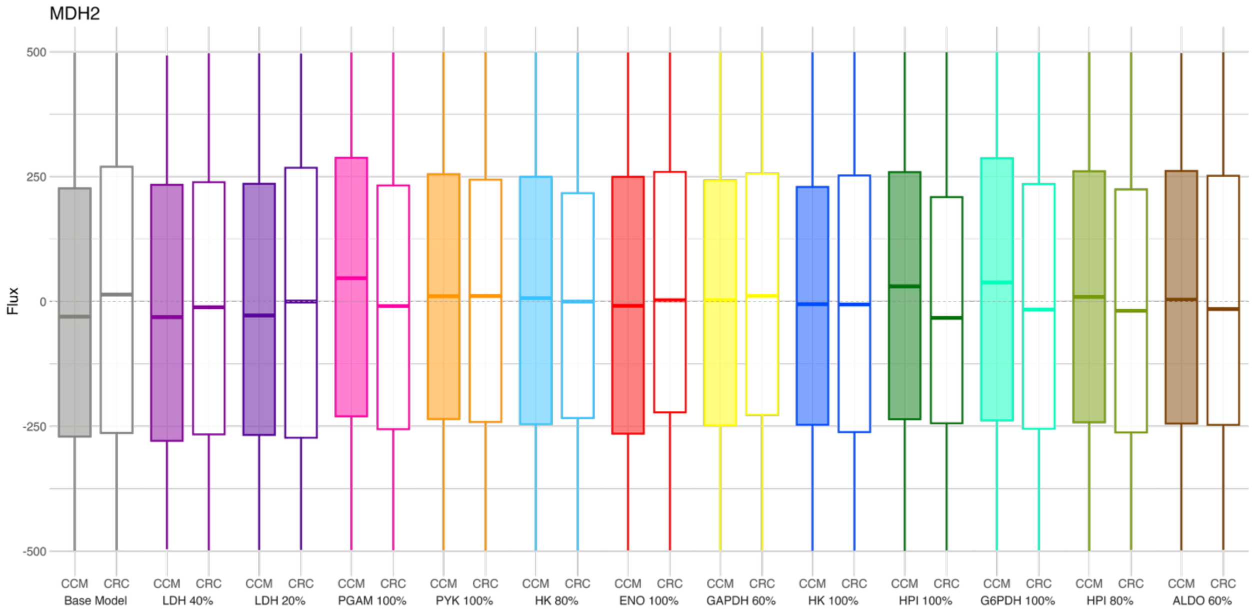
MDH2
Flux
500
250
0
-250
-500
CCM CRC
Base Model
LDH 40%
LDH 20%
PGAM 100%
PYK 100%
HK 80%
ENO 100%
GAPDH 60%
HK 100%
HPI 100%
G6PDH 100%
HPI 80%
ALDO 60%

FUM1

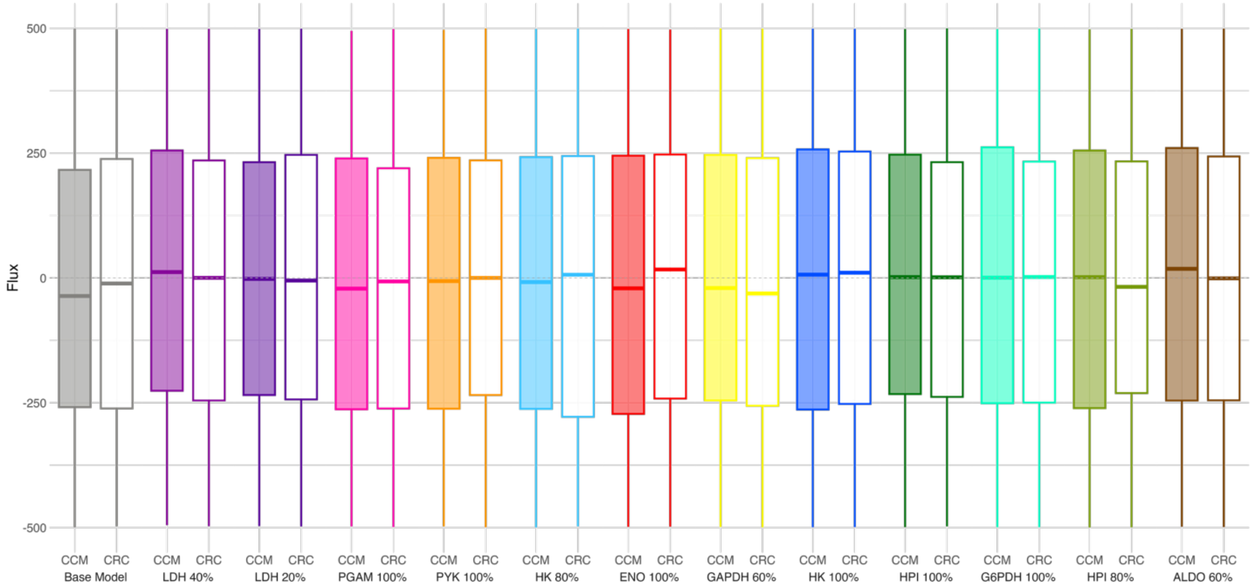

Flux
500
250
0
-250
-500
CCM CRC
Base Model
CCM CRC
LDH 40%
CCM CRC
LDH 20%
CCM CRC
PGAM 100%
CCM CRC
PYK 100%
CCM CRC
HK 80%
CCM CRC
ENO 100%
CCM CRC
GAPDH 60%
CCM CRC
HK 100%
CCM CRC
HPI 100%
CCM CRC
G6PDH 100%
CCM CRC
HPI 80%
CCM CRC
ALDO 60%


FUM2

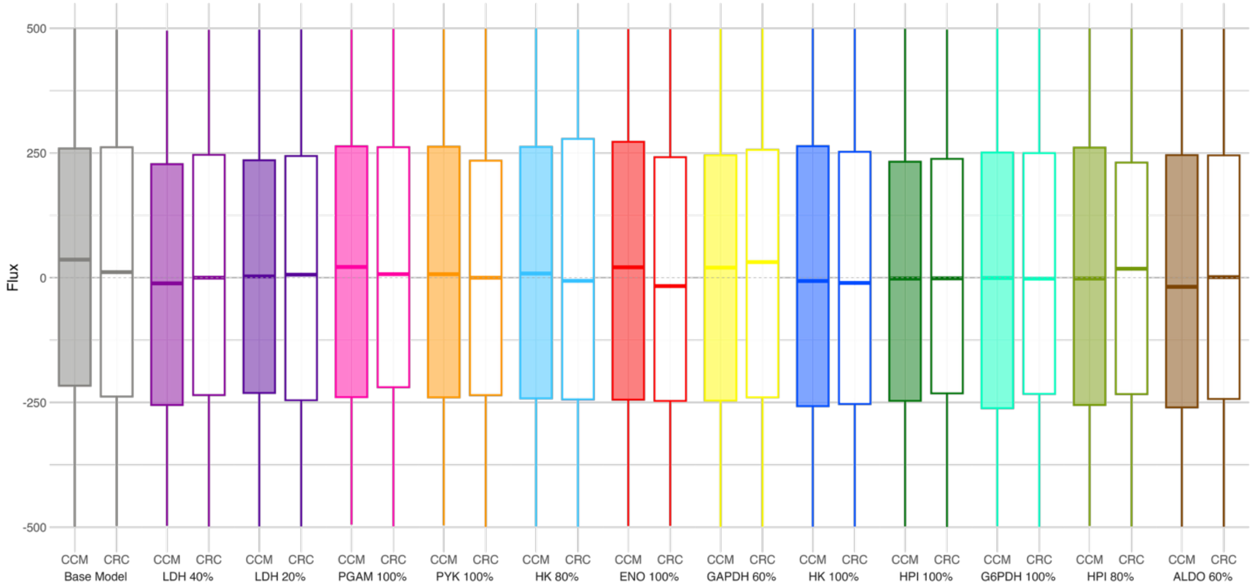

Flux
500
250
0
-250
-500
CCM CRC
Base Model
CCM CRC
LDH 40%
CCM CRC
LDH 20%
CCM CRC
PGAM 100%
CCM CRC
PYK 100%
CCM CRC
HK 80%
CCM CRC
ENO 100%
CCM CRC
GAPDH 60%
CCM CRC
HK 100%
CCM CRC
HPI 100%
CCM CRC
G6PDH 100%
CCM CRC
HPI 80%
CCM CRC
ALDO 60%

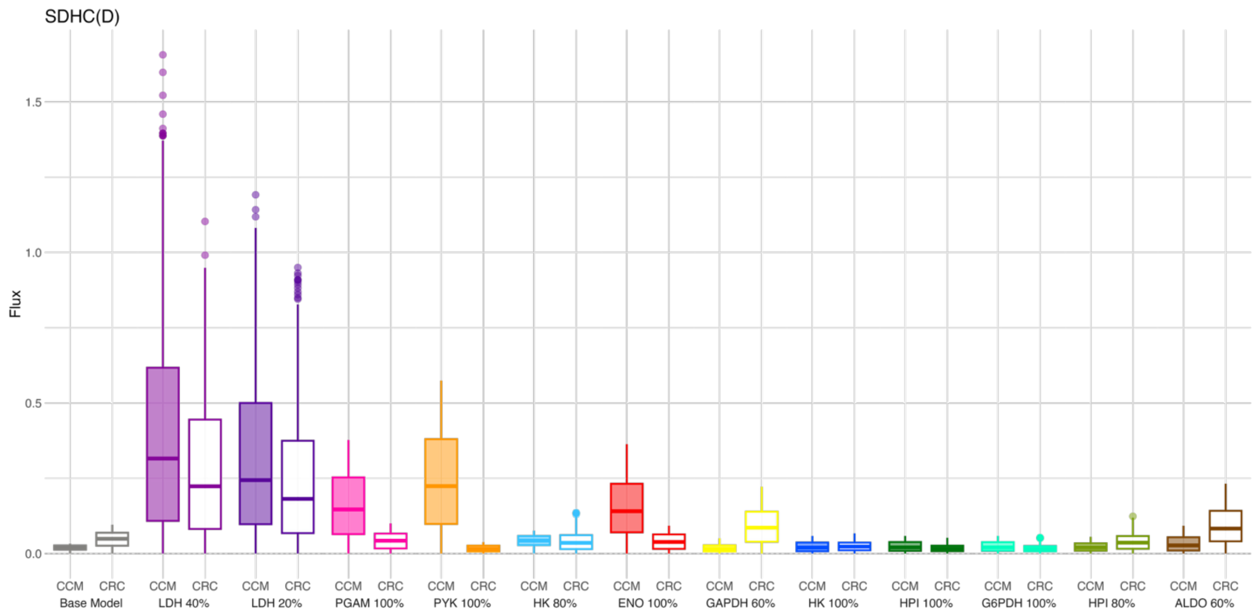
SDHC(D)
Flux
1.5
1.0
0.5
0.0
CCM CRC
Base Model
CCM CRC
LDH 40%
CCM CRC
LDH 20%
CCM CRC
PGAM 100%
CCM CRC
PYK 100%
CCM CRC
HK 80%
CCM CRC
ENO 100%
CCM CRC
GAPDH 60%
CCM CRC
HK 100%
CCM CRC
HPI 100%
CCM CRC
G6PDH 100%
CCM CRC
HPI 80%
CCM CRC
ALDO 60%

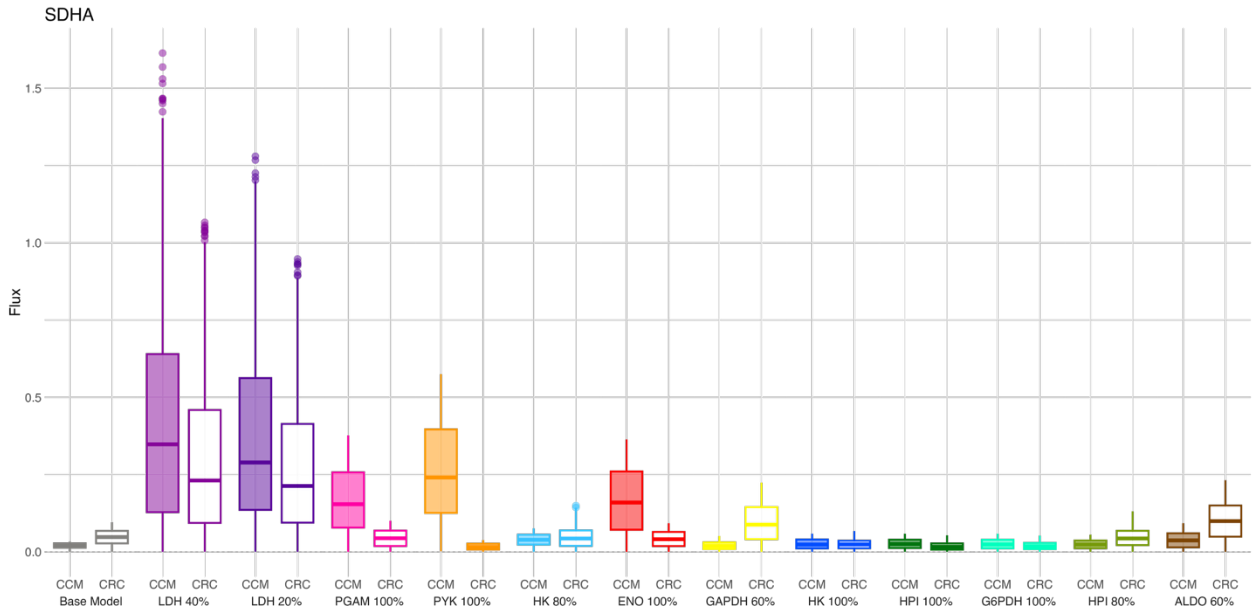
SDHA
Flux
1.5
1.0
0.5
0.0
CCM CRC
Base Model
CCM CRC
LDH 40%
CCM CRC
LDH 20%
CCM CRC
PGAM 100%
CCM CRC
PYK 100%
CCM CRC
HK 80%
CCM CRC
ENO 100%
CCM CRC
GAPDH 60%
CCM CRC
HK 100%
CCM CRC
HPI 100%
CCM CRC
G6PDH 100%
CCM CRC
HPI 80%
CCM CRC
ALDO 60%

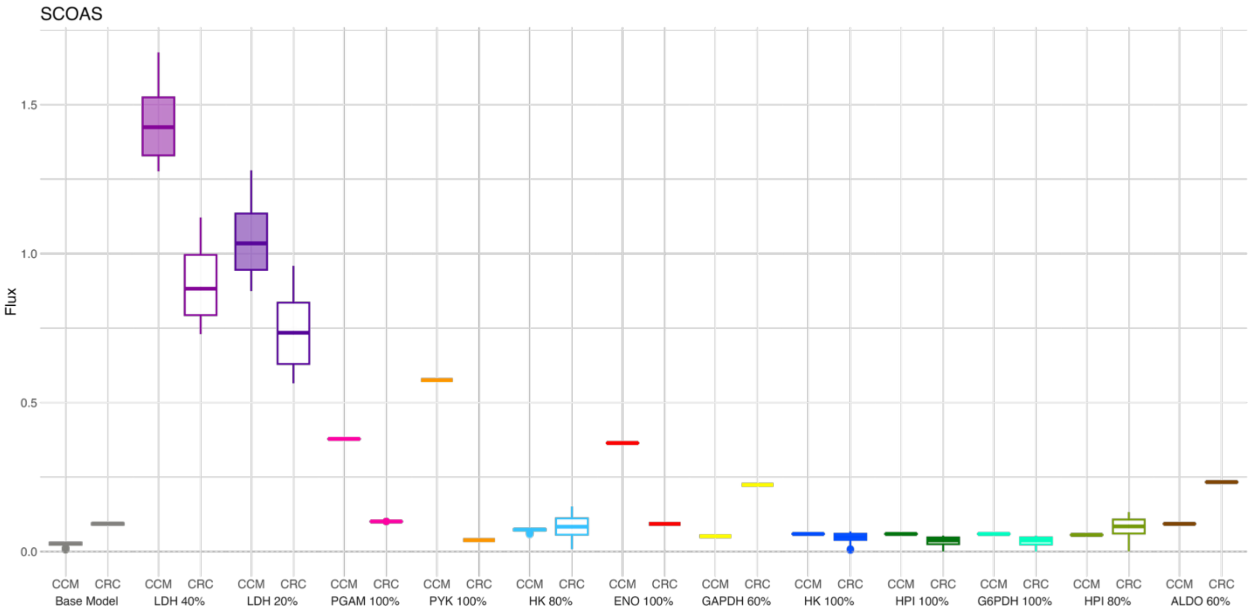

SCOAS
Flux
1.5
1.0
0.5
0.0
CCM CRC Base Model
CCM CRC LDH 40%
CCM CRC LDH 20%
CCM CRC PGAM 100%
CCM CRC PYK 100%
CCM CRC HK 80%
CCM CRC ENO 100%
CCM CRC GAPDH 60%
CCM CRC HK 100%
CCM CRC HPI 100%
CCM CRC G6PDH 100%
CCM CRC HPI 80%
CCM CRC ALDO 60%


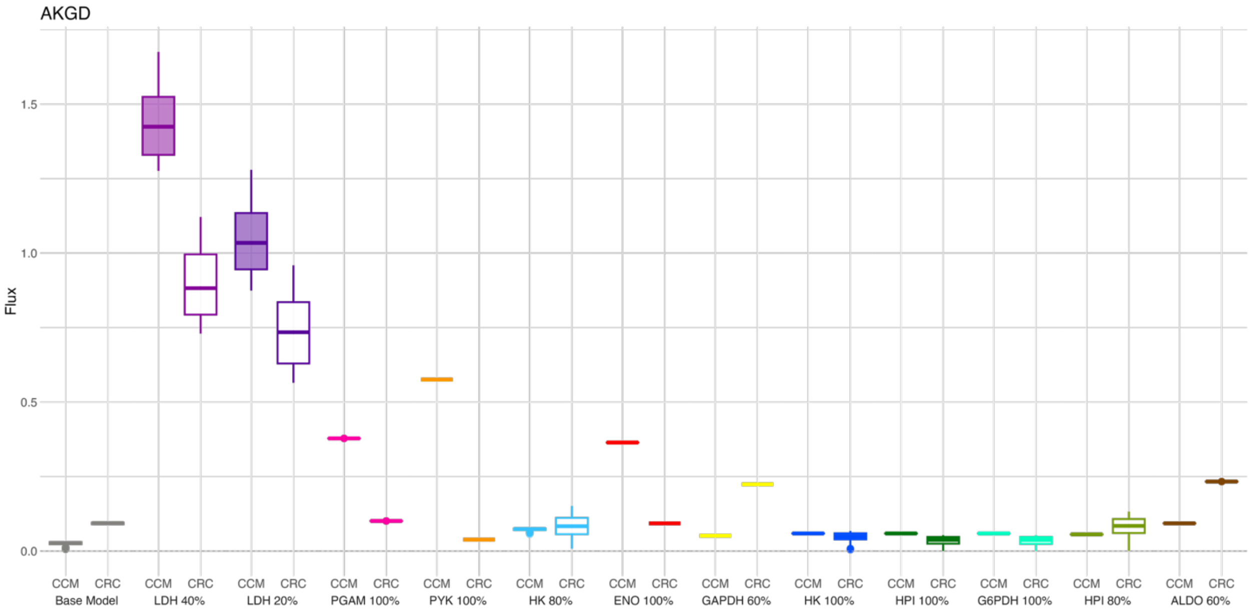

AKGD
Flux
1.5
1.0
0.5
0.0
CCM CRC Base Model
CCM CRC LDH 40%
CCM CRC LDH 20%
CCM CRC PGAM 100%
CCM CRC PYK 100%
CCM CRC HK 80%
CCM CRC ENO 100%
CCM CRC GAPDH 60%
CCM CRC HK 100%
CCM CRC HPI 100%
CCM CRC G6PDH 100%
CCM CRC HPI 80%
CCM CRC ALDO 60%

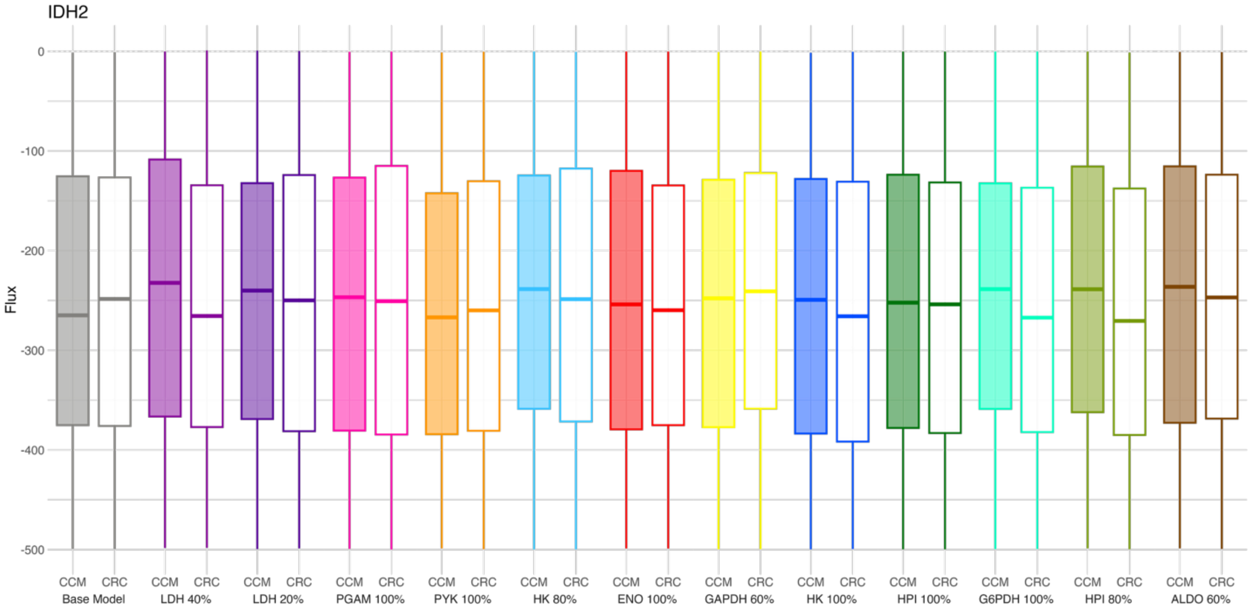
IDH2
Flux
0
-100
-200
-300
-400
-500
CCM CRC Base Model
CCM CRC LDH 40%
CCM CRC LDH 20%
CCM CRC PGAM 100%
CCM CRC PYK 100%
CCM CRC HK 80%
CCM CRC ENO 100%
CCM CRC GAPDH 60%
CCM CRC HK 100%
CCM CRC HPI 100%
CCM CRC G6PDH 100%
CCM CRC HPI 80%
CCM CRC ALDO 60%

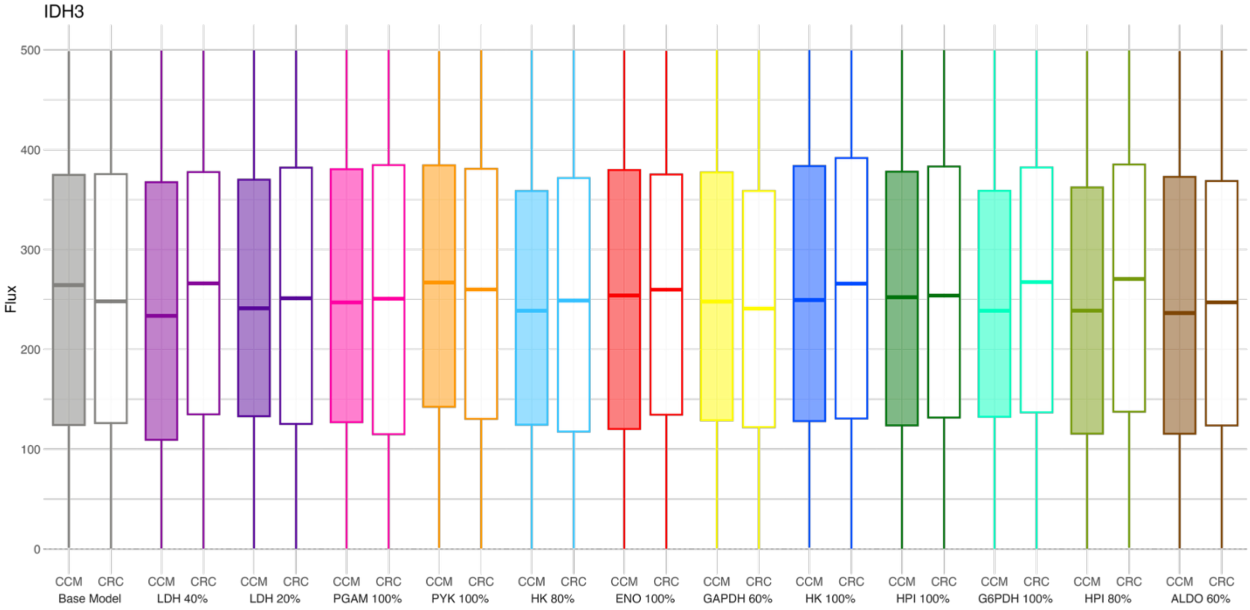
IDH3
Flux
500
400
300
200
100
0
CCM CRC Base Model
CCM CRC LDH 40%
CCM CRC LDH 20%
CCM CRC PGAM 100%
CCM CRC PYK 100%
CCM CRC HK 80%
CCM CRC ENO 100%
CCM CRC GAPDH 60%
CCM CRC HK 100%
CCM CRC HPI 100%
CCM CRC G6PDH 100%
CCM CRC HPI 80%
CCM CRC ALDO 60%

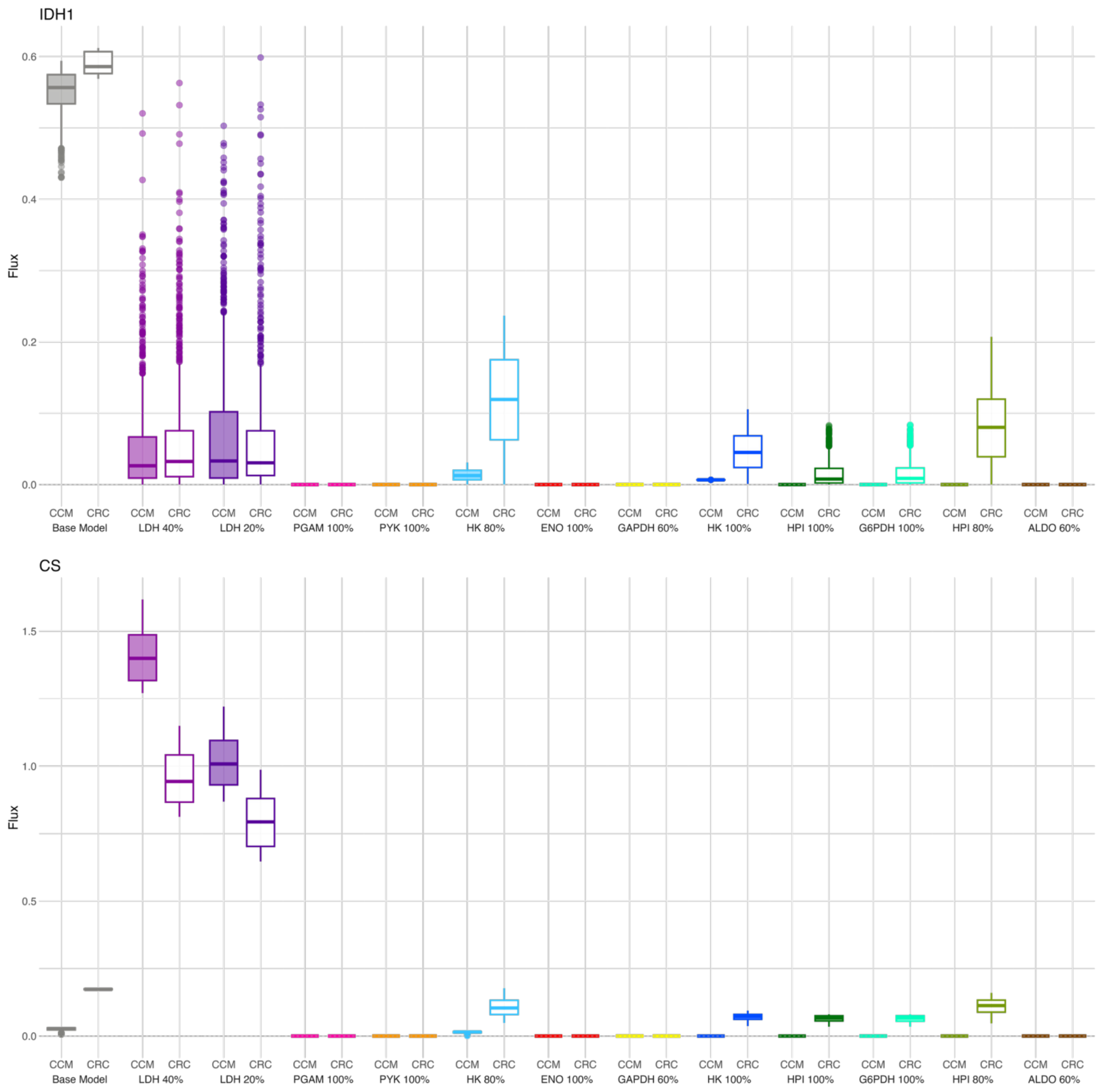
IDH1
Flux
0.0
0.2
0.4
0.6
CCM CRC
Base Model
LDH 40%
LDH 20%
PGAM 100%
PYK 100%
HK 80%
ENO 100%
GAPDH 60%
HK 100%
HPI 100%
G6PDH 100%
HPI 80%
ALDO 60%
CS
Flux
0.0
0.5
1.0
1.5
CCM CRC
Base Model
LDH 40%
LDH 20%
PGAM 100%
PYK 100%
HK 80%
ENO 100%
GAPDH 60%
HK 100%
HPI 100%
G6PDH 100%
HPI 80%
ALDO 60%

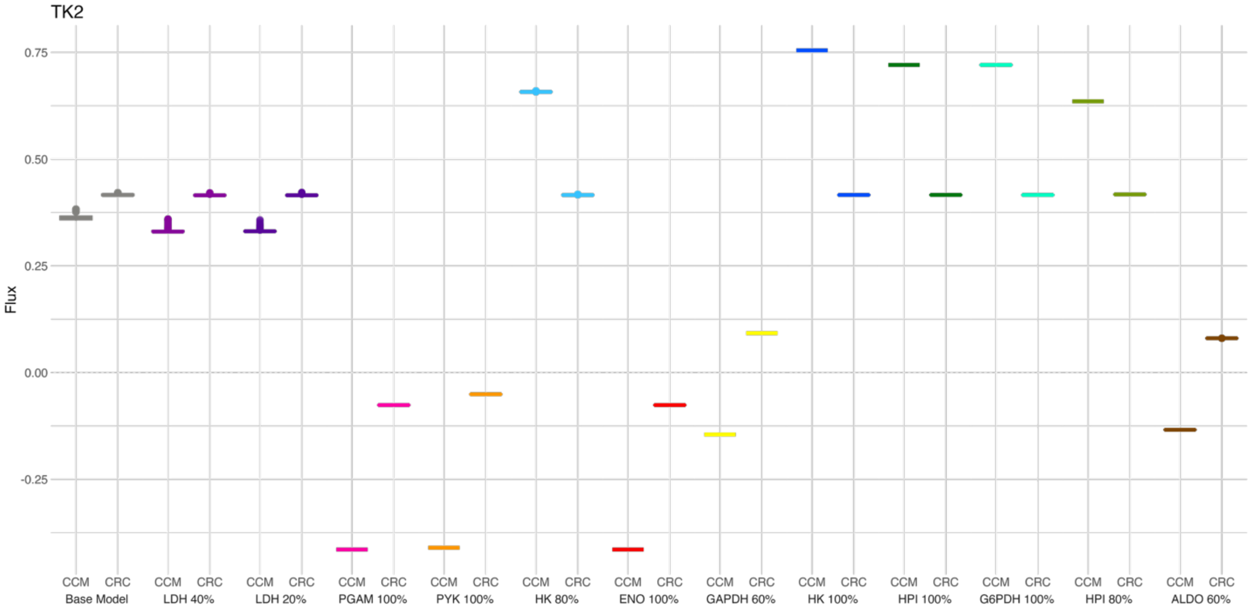
TK2
0.75
0.50
0.25
0.00
-0.25
Flux
CCM CRC
Base Model
LDH 40%
LDH 20%
PGAM 100%
PYK 100%
HK 80%
ENO 100%
GAPDH 60%
HK 100%
HPI 100%
G6PDH 100%
HPI 80%
ALDO 60%

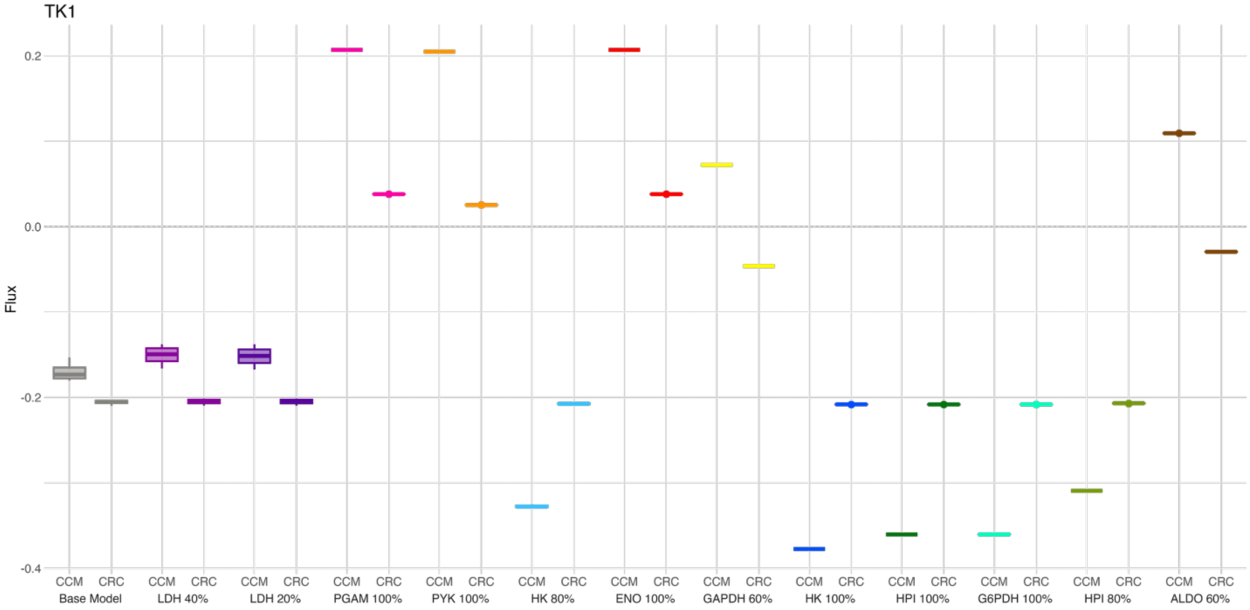
TK1
0.2
0.0
-0.2
-0.4
Flux
CCM CRC
Base Model
LDH 40%
LDH 20%
PGAM 100%
PYK 100%
HK 80%
ENO 100%
GAPDH 60%
HK 100%
HPI 100%
G6PDH 100%
HPI 80%
ALDO 60%

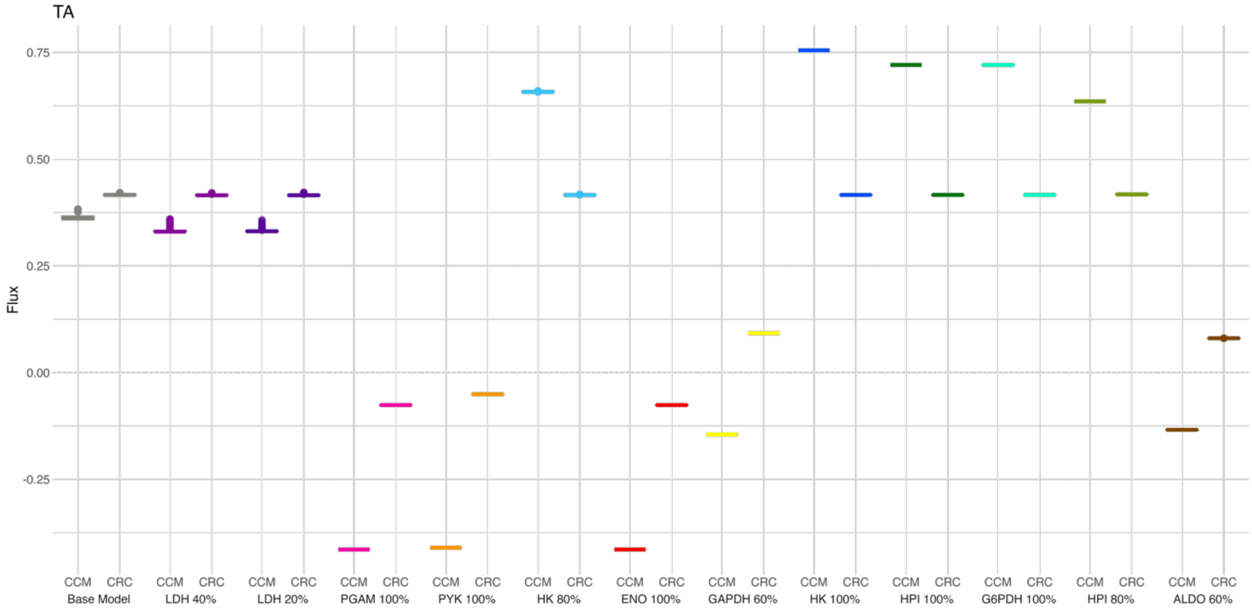
TA
Flux
0.75
0.50
0.25
0.00
-0.25
CCM CRC
Base Model
LDH 40%
LDH 20%
PGAM 100%
PYK 100%
HK 80%
ENO 100%
GAPDH 60%
HK 100%
HPI 100%
G6PDH 100%
HPI 80%
ALDO 60%

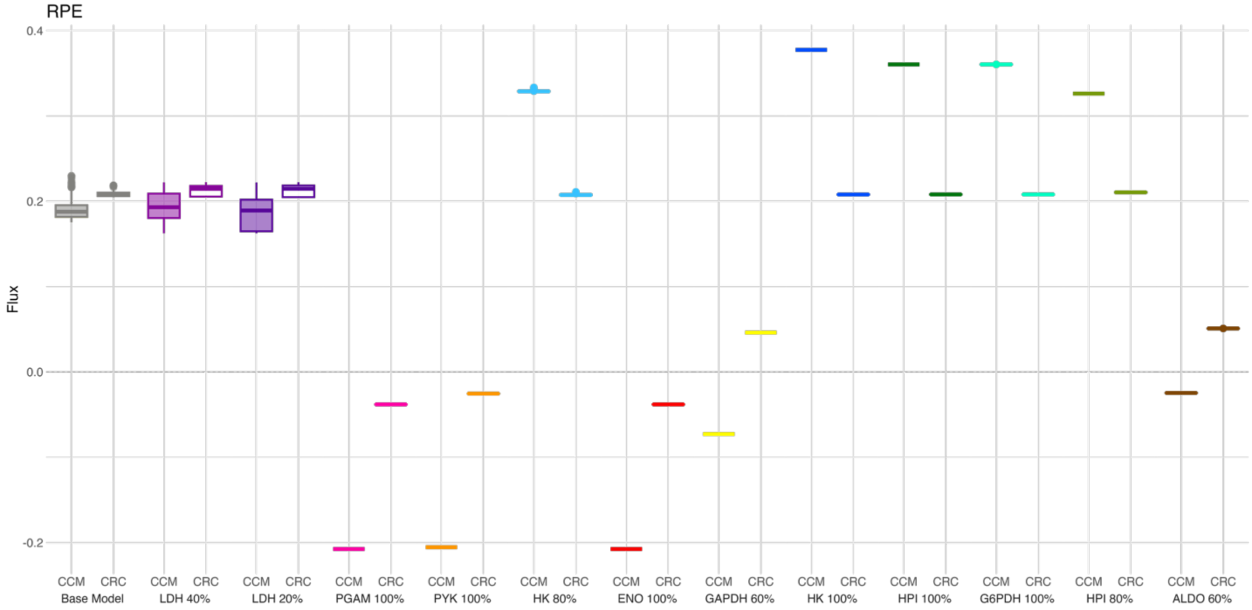
RPE
Flux
0.4
0.2
0.0
-0.2
CCM CRC
Base Model
LDH 40%
LDH 20%
PGAM 100%
PYK 100%
HK 80%
ENO 100%
GAPDH 60%
HK 100%
HPI 100%
G6PDH 100%
HPI 80%
ALDO 60%

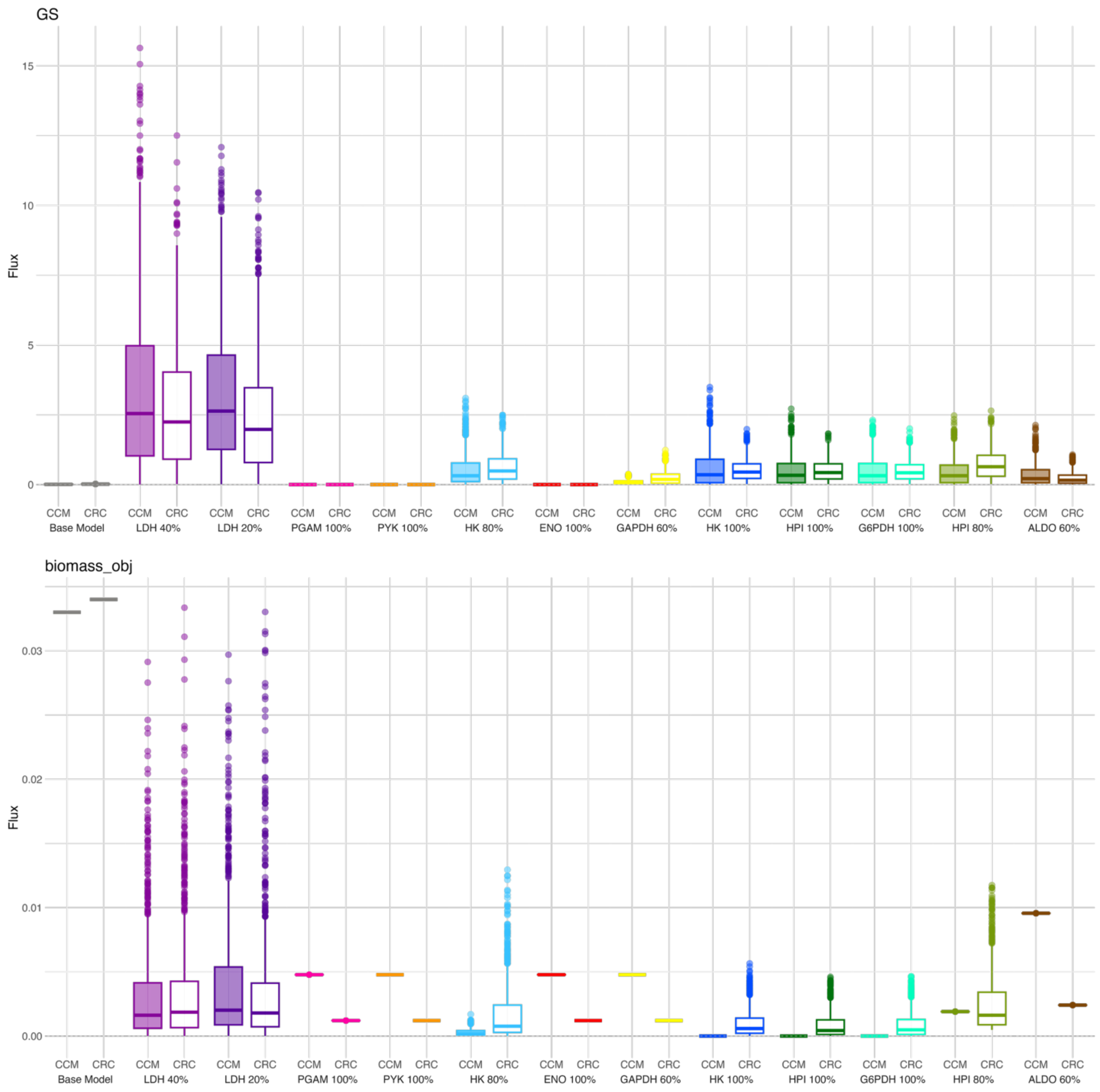

GS
Flux
15
10
5
0
CCM CRC Base Model
CCM CRC LDH 40%
CCM CRC LDH 20%
CCM CRC PGAM 100%
CCM CRC PYK 100%
CCM CRC HK 80%
CCM CRC ENO 100%
CCM CRC GAPDH 60%
CCM CRC HK 100%
CCM CRC HPI 100%
CCM CRC G6PDH 100%
CCM CRC HPI 80%
CCM CRC ALDO 60%
biomass_obj
Flux
0.03
0.02
0.01
0.00
CCM CRC Base Model
CCM CRC LDH 40%
CCM CRC LDH 20%
CCM CRC PGAM 100%
CCM CRC PYK 100%
CCM CRC HK 80%
CCM CRC ENO 100%
CCM CRC GAPDH 60%
CCM CRC HK 100%
CCM CRC HPI 100%
CCM CRC G6PDH 100%
CCM CRC HPI 80%
CCM CRC ALDO 60%

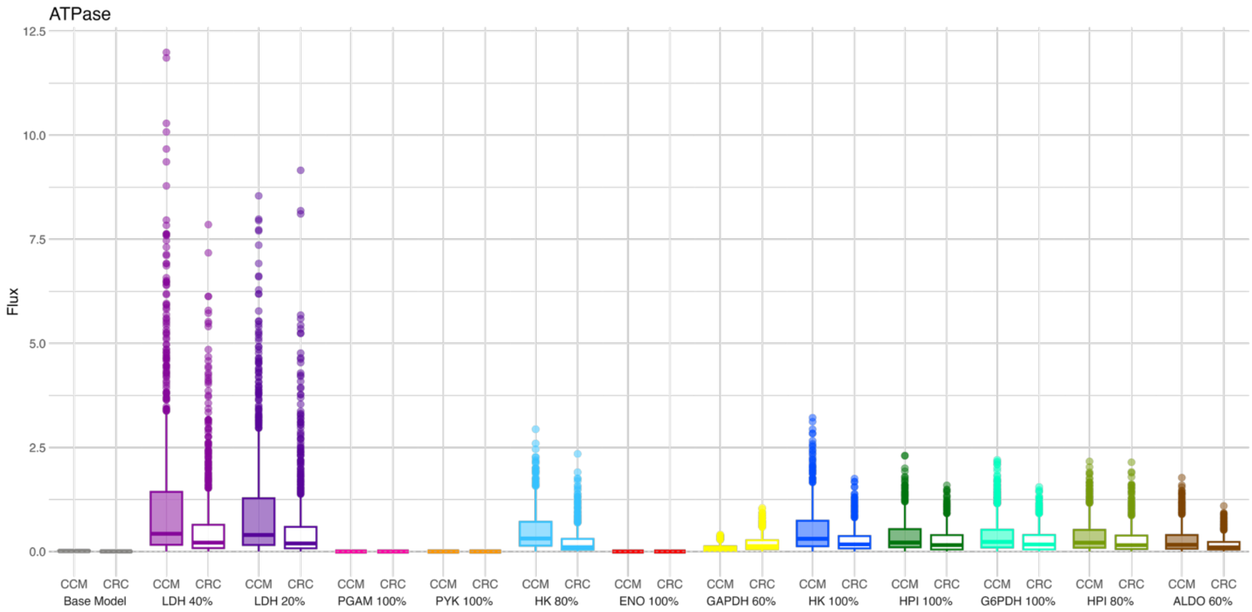

ATPase
Flux
12.5
10.0
7.5
5.0
2.5
0.0
CCM CRC
Base Model
LDH 40%
LDH 20%
PGAM 100%
PYK 100%
HK 80%
ENO 100%
GAPDH 60%
HK 100%
HPI 100%
G6PDH 100%
HPI 80%
ALDO 60%


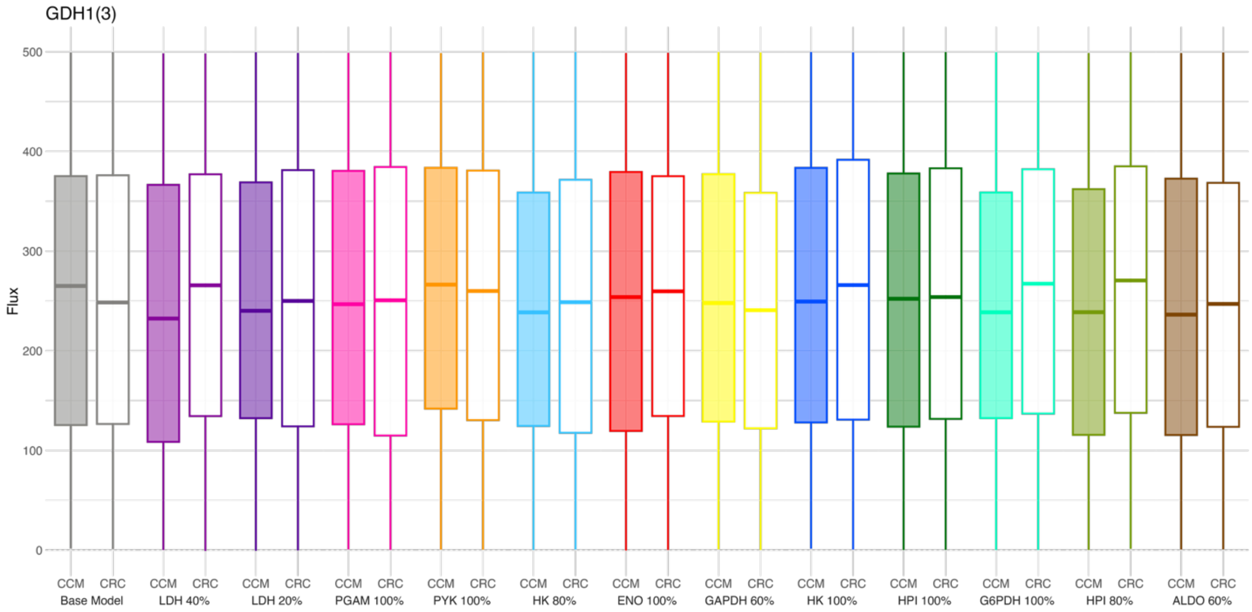

GDH1(3)
Flux
500
400
300
200
100
0
CCM CRC
Base Model
LDH 40%
LDH 20%
PGAM 100%
PYK 100%
HK 80%
ENO 100%
GAPDH 60%
HK 100%
HPI 100%
G6PDH 100%
HPI 80%
ALDO 60%

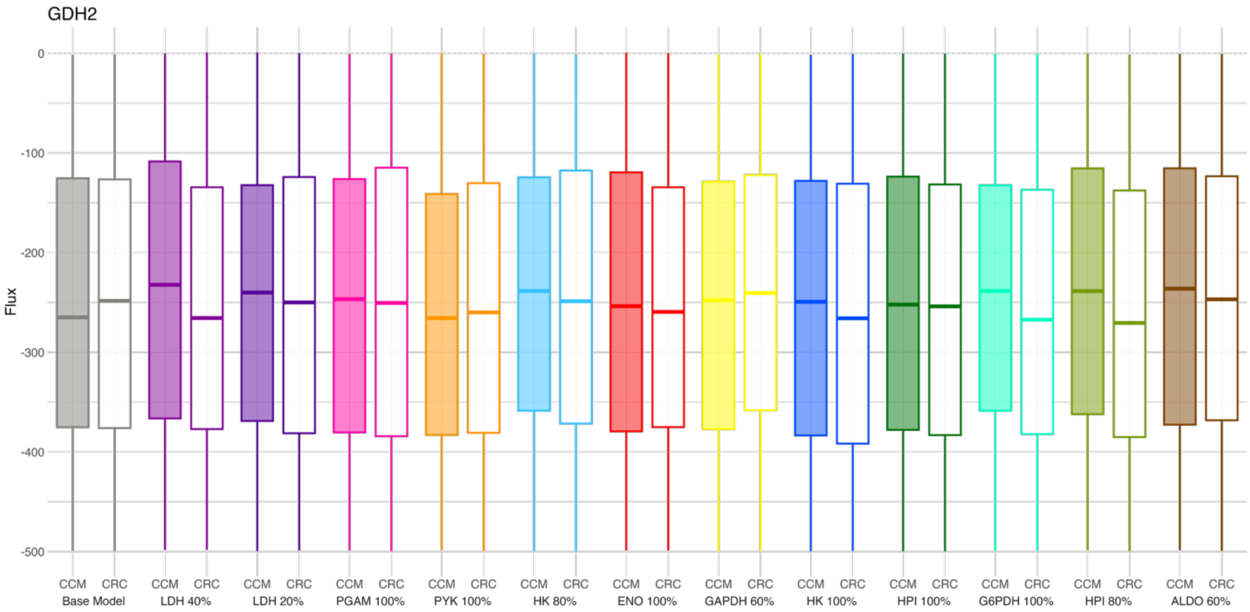
GDH2
Flux
0
-100
-200
-300
-400
-500
CCM CRC Base Model
CCM CRC LDH 40%
CCM CRC LDH 20%
CCM CRC PGAM 100%
CCM CRC PYK 100%
CCM CRC HK 80%
CCM CRC ENO 100%
CCM CRC GAPDH 60%
CCM CRC HK 100%
CCM CRC HPI 100%
CCM CRC G6PDH 100%
CCM CRC HPI 80%
CCM CRC ALDO 60%

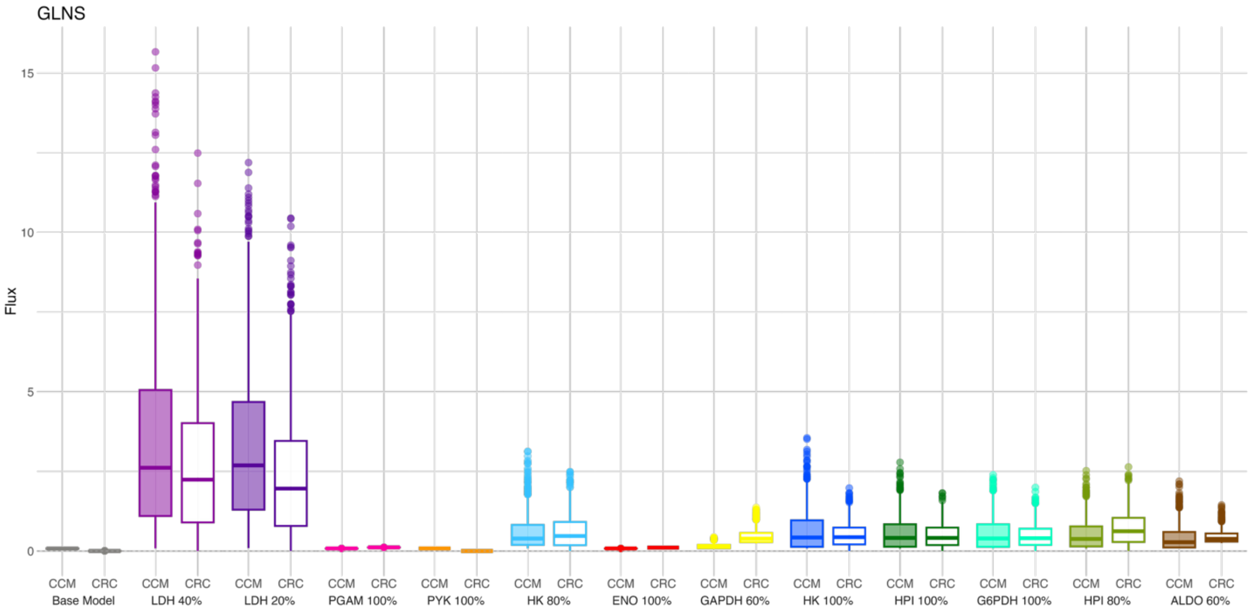
GLNS
Flux
15
10
5
0
CCM CRC Base Model
CCM CRC LDH 40%
CCM CRC LDH 20%
CCM CRC PGAM 100%
CCM CRC PYK 100%
CCM CRC HK 80%
CCM CRC ENO 100%
CCM CRC GAPDH 60%
CCM CRC HK 100%
CCM CRC HPI 100%
CCM CRC G6PDH 100%
CCM CRC HPI 80%
CCM CRC ALDO 60%

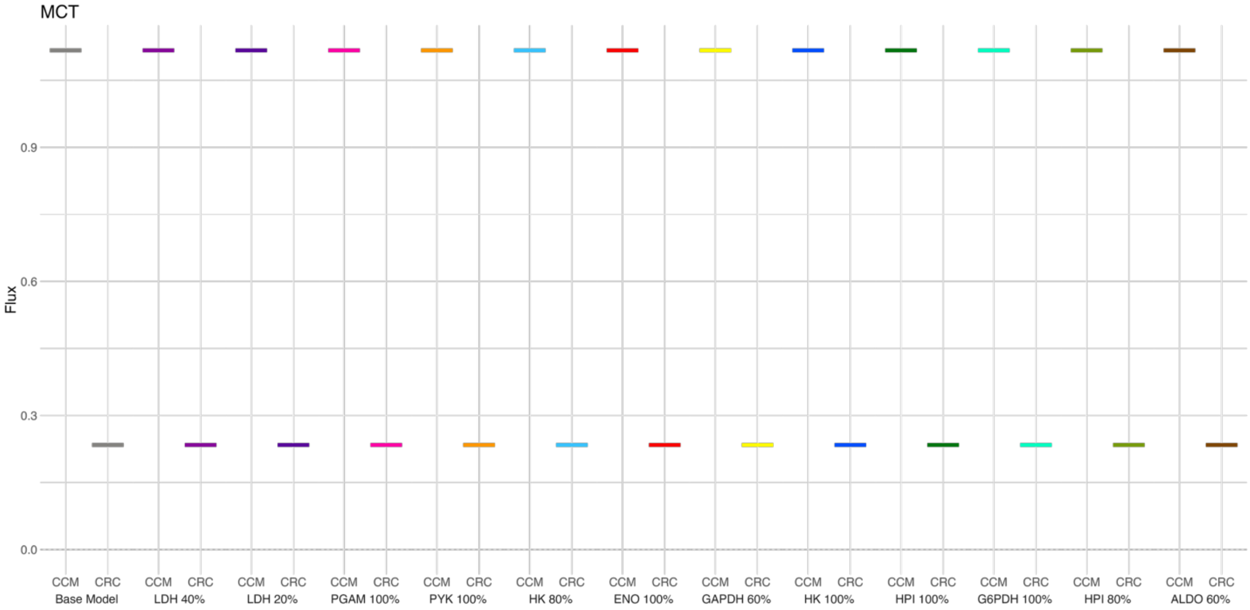
MCT
Flux
0.0
0.3
0.6
0.9
CCM CRC Base Model
CCM CRC LDH 40%
CCM CRC LDH 20%
CCM CRC PGAM 100%
CCM CRC PYK 100%
CCM CRC HK 80%
CCM CRC ENO 100%
CCM CRC GAPDH 60%
CCM CRC HK 100%
CCM CRC HPI 100%
CCM CRC G6PDH 100%
CCM CRC HPI 80%
CCM CRC ALDO 60%

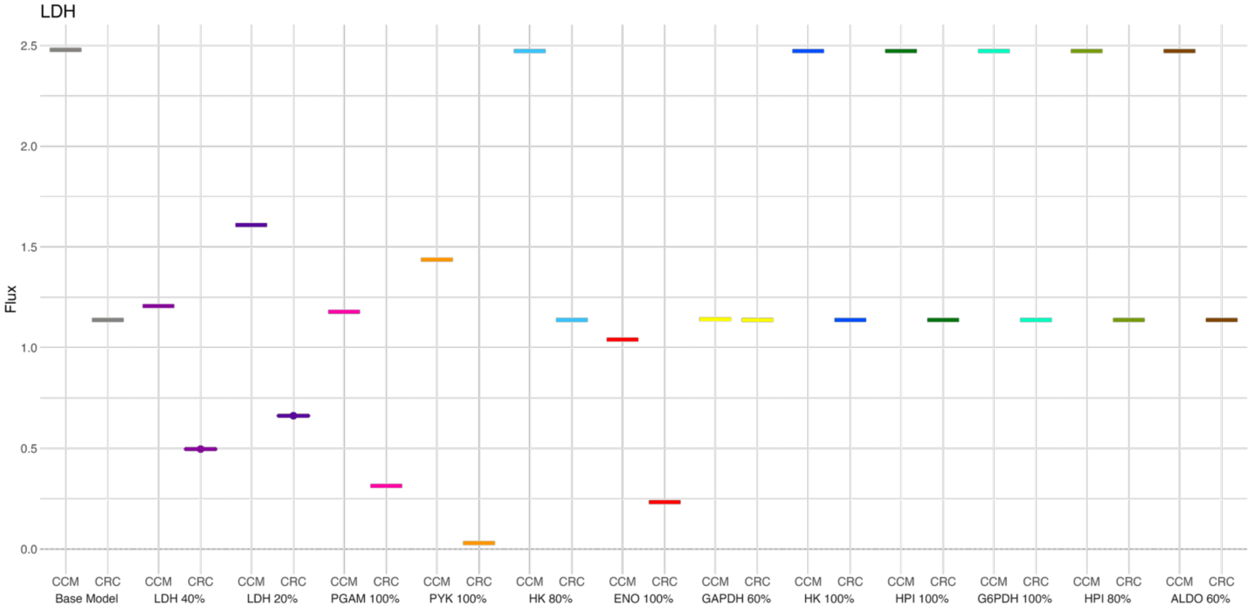
LDH
Flux
0.0
0.5
1.0
1.5
2.0
2.5
CCM CRC Base Model
CCM CRC LDH 40%
CCM CRC LDH 20%
CCM CRC PGAM 100%
CCM CRC PYK 100%
CCM CRC HK 80%
CCM CRC ENO 100%
CCM CRC GAPDH 60%
CCM CRC HK 100%
CCM CRC HPI 100%
CCM CRC G6PDH 100%
CCM CRC HPI 80%
CCM CRC ALDO 60%

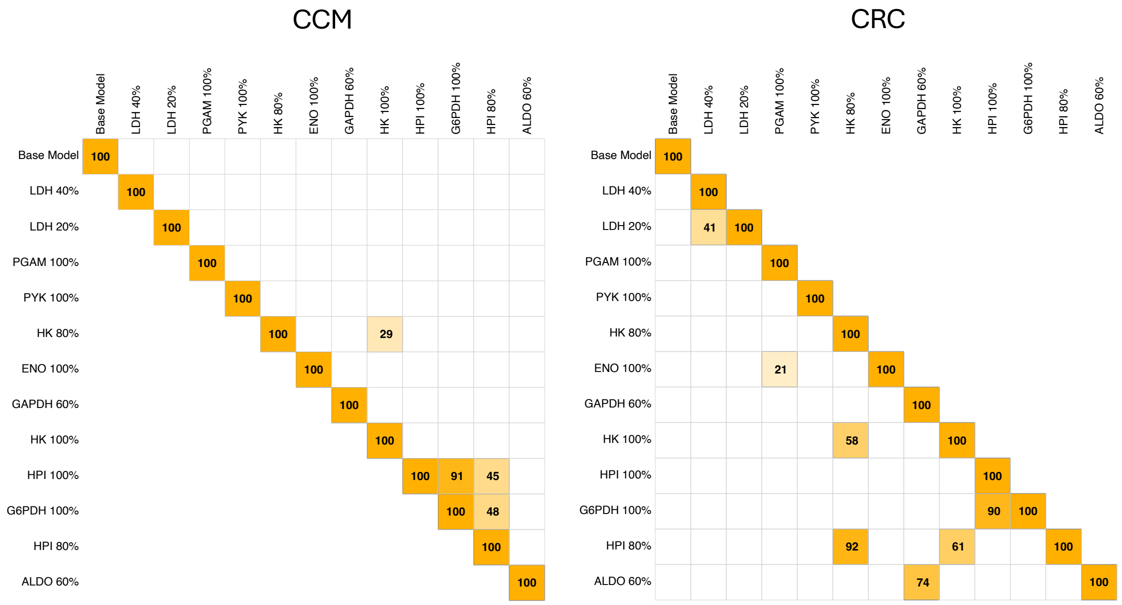


**Supplemental Figure 2. Percent overlap between knockdown model clusters indicates shared metabolic state.** Quantification of shared metabolic states is based on the percent overlap between clusters. Percent overlap indicated by shade, where darker orange represents greater overlap, and by value inside each cell.

| | Mean Distance From Point to Centroid | | Distance Between Centroids | | |
|---|---|---|---|---|---|
| **Model** | **CCM** | **CRC** | **CCM & Base** | **CRC & Base** | **CCM & CRC** |
| **Base Model** | 1.48 ± 0.82 | 1.28 ± 0.73 | 0.00 | 0.00 | 24.53 |
| **ALDO 60%** | 1.41 ± 0.76 | 1.07 ± 0.56 | 25.14 | 26.57 | 44.32 |
| **ENO 100%** | 1.07 ± 0.59 | 1.12 ± 0.62 | 54.87 | 40.46 | 14.65 |
| **G6PDH 100%** | 1.06 ± 0.59 | 1.08 ± 0.59 | 19.54 | 16.24 | 18.98 |
| **GAPDH 60%** | 1.20 ± 0.64 | 1.05 ± 0.57 | 40.35 | 26.04 | 11.27 |
| **HK 100%** | 1.14 ± 0.66 | 1.13 ± 0.61 | 14.64 | 12.10 | 40.91 |
| **HK 80%** | 1.12 ± 0.66 | 1.18 ± 0.65 | 11.65 | 10.16 | 36.77 |
| **HPI 100%** | 1.06 ± 0.57 | 1.06 ± 0.55 | 19.50 | 16.25 | 19.02 |
| **HPI 80%** | 1.10 ± 0.60 | 1.20 ± 0.65 | 17.08 | 10.26 | 14.20 |
| **LDH 20%** | 1.65 ± 0.87 | 1.42 ± 0.78 | 23.34 | 13.93 | 21.24 |
| **LDH 40%** | 1.63 ± 0.90 | 1.41 ± 0.75 | 34.27 | 18.32 | 20.74 |
| **PGAM 100%** | 1.10 ± 0.58 | 1.10 ± 0.60 | 50.48 | 38.43 | 17.92 |
| **PYK 100%** | 1.12 ± 0.61 | 1.17 ± 0.65 | 65.75 | 47.41 | 19.42 |

**Supplemental Table 1.** Distance between flux sampling points in representation learning projection. Distance between centroids presented as mean, and distance from point to centroid presented as mean and standard deviation.

| | Predicted Biomass Flux | |
|---|---|---|
| **Model** | **CCM** | **CRC** |
| **Base Model** | $3.3x10^{-2}$ | $3.4 \times 10^{-2}$ |
| **ALDO 60%** | $9.56x10^{-3}$ | $2.41x10^{-3}$ |
| **ENO 100%** | $4.78x10^{-3}$ | $1.21x10^{-3}$ |
| **G6PDH 100%** | - | $8.58x10^{-4} \pm 9.51x10^{-4}$ |
| **GAPDH 60%** | $4.78x10^{-3}$ | $1.21x10^{-3}$ |
| **HK 80%** | $22.87x10^{-4} \pm 2.76x10^{-4}$ | $1.77x10^{-3} \pm 2.25x10^{-3}$ |
| **HK 100%** | - | $9.69x10^{-4} \pm 1.03x10^{-3}$ |
| **HPI 80%** | $1.9x10^{-3}$ | $2.55x10^{-3} \pm 2.32x10^{-3}$ |
| **HPI 100%** | - | $8.36x10^{-4} \pm 9.65x10^{-4}$ |
| **LDH 20%** | $3.99x10^{-3} \pm 4.84x10^{-3}$ | $3.42x10^{-3} \pm 4.74x10^{-3}$ |
| **LDH 40%** | $3.2x10^{-3} \pm 4.14x10^{-3}$ | $3.47x10^{-3} \pm 4.48x10^{-3}$ |
| **PGAM 100%** | $44.78x10^{-3}$ | $1.21x10^{-3}$ |
| **PYK 100%** | $4.78x10^{-3}$ | $1.21x10^{-3}$ |
| **Knockdown Average** | $3.17x10^{-3} \pm 2.89x10^{-3}$ | $1.76x10^{-3} \pm 9.66x10^{-4}$ |

**Supplemental Table 2.** Mean and standard deviation predicted biomass flux.
